# Deciphering the RNA-binding protein network during endosomal mRNA transport

**DOI:** 10.1101/2024.03.22.586338

**Authors:** Senthil-Kumar Devan, Sainath Shanmugasundaram, Kira Müntjes, Sander HJ Smits, Florian Altegoer, Michael Feldbrügge

## Abstract

Microtubule-dependent endosomal transport is crucial for polar growth, ensuring the precise distribution of cellular cargos such as proteins and mRNAs. However, the molecular mechanism linking mRNAs to the endosomal surface remains poorly understood. Here, we present a structural analysis of the key RNA-binding protein Rrm4 from *Ustilago maydis*. Our findings reveal a new type of MademoiseLLE domain featuring a seven-helical bundle that provides a distinct binding interface. A comparative analysis with the canonical MLLE domain of the poly(A)-binding protein Pab1 disclosed unique characteristics of both domains. Deciphering the MLLE binding code enabled prediction and verification of previously unknown Rrm4 interactors containing short linear motifs. Importantly, we demonstrated that the human MLLE domains, such as those of PABPC1 and UBR5, employed a similar principle to distinguish among interaction partners. Thus, our study provides unprecedented mechanistic insights into how structural variations in the widely distributed MLLE domain facilitates mRNA attachment during endosomal transport.

**Significance:** Polar growing cells, such as fungal hyphae and neurons, utilize endosomes to transport mRNAs along their microtubules. But how do these mRNAs precisely attach to endosomes? Our study addresses this question by investing the key mRNA transporter, Rrm4, in a fungal model microorganism. We uncovered new features of a protein-protein interaction domain that recognizes specific short linear motifs in binding partners. While this domain resembles one found in the poly(A)-binding protein, it exhibits distinct motif recognition. Deciphering the underlying binding code unveiled new interaction partners for Rrm4. The recognition system is used to form a resilient network of RNA-binding proteins (RBPs) and their interaction partners during endosomal transport. This principle is applicable to humans, highlighting its fundamental importance.

## Introduction

Highly polarized cells, such as fungal hyphae, depend on active long-distance transport along the cytoskeleton to sustain efficient expansion at the growth pole (1). This dependency is particularly evident in pathogens that employ efficient hyphal growth as part of their infection strategy (2, 3). Microtubule-dependent trafficking is facilitated by Rab5a-positive endosomes, which transports various cargos, including proteins, lipids, mRNA, ribosomes and even entire organelles such as peroxisomes (4–6). Endosomes shuttle throughout the hyphae via the concerted action of plus-end directed kinesin and minus-end directed dynein motors along microtubules (7, 8). However, detailed mechanistic insights into how protein-protein interactions govern cargo specificity and endosomal attachment, are still lacking.

We study the fungal pathogen *Ustilago maydis,* which causes smut disease in maize. Pathogenicity relies on the formation of infectious hyphae, which are reliant on microtubule-dependent endosomal mRNA transport (5, 9). The key RNA-binding protein Rrm4 plays a pivotal role by binding to hundreds of mRNA targets, facilitating the distribution of mRNAs and associated ribosomes throughout the hyphae (10–12). Notably, cargo mRNAs encoding all four septins undergo endosome-coupled translation, which is essential for the assembly of heteromeric septin complexes on the surface of endosomes. These complexes are subsequently transported toward the hyphal tip to form higher-septin filaments (10, 13).

Rrm4 comprises three N-terminal RNA recognition motifs (RRMs) for cargo recognition and three C-terminal MademoiseLLE (MLLE) domains for protein-protein interactions (Fig. 1A; 14). The MLLE domains serve as a binding platform with a distinct hierarchy: while the first and second MLLE domains (MLLE1^Rrm4^, MLLE2^Rrm4^) fulfil accessory functions, the third MLLE domain (MLLE3^Rrm4^) is crucial for interaction with the two PAM2-like sequences (PAM2L) of the adaptor protein Upa1, a FYVE zinc finger-containing protein that is essential for endosomal Rrm4 attachment (Fig 1A; 14, 15). Loss of Upa1 results in severe defects in endosomal shuttling of Rrm4-containing mRNAs. However, residual shuttling of Rrm4 is still observed, suggesting the presence of at least one additional, currently unknown, adaptor protein.

**Figure 1.**
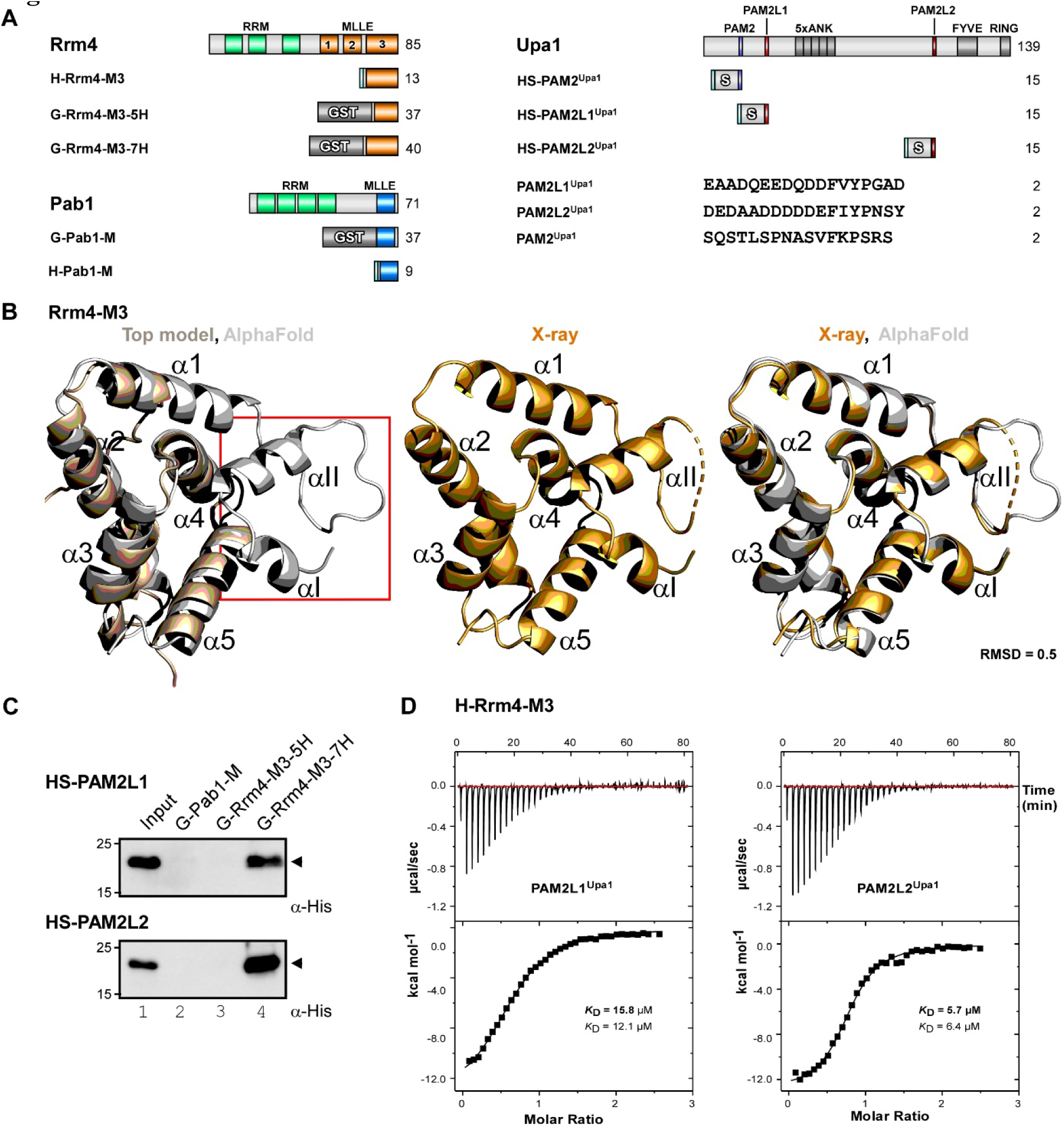
A seven-helix type MLLE domain confers specific binding. (**A**) Schematic representation of protein variants (molecular weight in kilo Dalton, green, RNA recognition motif (RRM); orange, MLLE^Rrm4^ domains; bright blue, MLLE^Pab1^; light blue PAM2^Upa1^; light red PAM2L1^Upa1^; dark red PAM2L2^Upa1^; dark grey, Ankyrin repeats (5xANK), FYVE domain, and RING domain of Upa1, cyan, His_6_. Sequences of PAM2 and PAM2L1,2 peptides are denoted. The following symbols are used: M3, MLLE3^Rrm4^; M, MLLE^Pab1^; 5H, five helices; 7H, seven helices, G, GST tag; HS, His_6_-Sumo tag. (**B**) 3D structural models of MLLE3^Rrm4^ domain, generated using TopModel, Alphafold, and X-ray as indicated. (**C**) Western blot analysis of GST pull-down experiments using α-His for detection (input, respective His_6_-SUMO peptides). (**D**) Representative isothermal titration calorimetry (ITC) binding curves of MLLE3^Rrm4^ domain (H-Rrm4-M3). *K*_D_ values of two independent measurements are given (indicated data in bold).

The MLLE domain was initially discovered in cytoplasmic poly(A)-binding protein, which binds to most poly(A) tails in eukaryotic mRNAs (PABPC1, Pab1p and Pab1 in *H. sapiens*, *S. cerevisiae* and *U. maydis*, respectively; 16-18). The MLLE domain of PABPC1 (MLLE^PABPC1^), has been shown to interact with PAM2 motifs of numerous interaction partners, such as GW182 and eRF3, which function in microRNA biology and translational termination, respectively (19). PAM2 serves as a prime example of a short linear motif (SLiM). SLiMs are widespread recognition motifs found in unstructured regions of RNA-binding proteins or their adaptors, enabling the formation of higher-order interaction networks. It has been proposed, for example, that 64% of the 120 PABPC1 interaction partners harbor a PAM2 or sequence similar SLiM sequences (20, 21).

In MLLE^PABPC1^, the core structure is consisted of a five α-helical bundle. In the classical binding mode, the PAM2 motif binds to the MLLE by interacting with hydrophobic pockets between helices 2-3 and 3-5. This interaction is facilitated by the highly conserved phenylalanine and leucine residues of PAM2 (19, 22). Currently, only one exception is known: the PAM2 sequences of GW182 binds to a different interface along helix 2 (23). A second MLLE domain is present in UBR5, a HECT-type E3 ubiquitin ligase functioning as chain-elongating E3 ubiquitin ligase (<u>H</u>omologous to <u>E</u>6AP <u>C</u>-<u>T</u>erminus; 24). This domain interacts with PAM2 sequences such as PAM2^PAIP^, with high affinity (25). However, the biological function of its MLLE domain is currently unclear (26–28).

The endosomal adaptor protein Upa1 also contains a PAM2 motif (PAM2^Upa1^) for interaction with the MLLE domain of Pab1 (MLLE^Pab1^, Fig. 1A; 15). Furthermore, an essential scaffold protein of endosomal mRNAs, Upa2, also harbors four PAM2 sequences. Notably, the MLLE^Pab1^ and MLLE3^Rrm4^ exhibit an exquisite specificity in differentiating between PAM2 and PAM2L sequences (14). In essence, endosomal messenger ribonucleoproteins (mRNPs) containing the MLLE domain proteins Rrm4 and Pab1 are attached to endosomes through interaction with PAM2 and PAM2L sequences in a complex manner. To understand the underlying SLiM binding code, we aimed to clarify specific recognition of MLLE domains and their cognate PAM2 or PAM2L sequences at the structural level. We disclose a new domain architecture of MLLE3^Rrm4^ and uncover mechanistic details that enabled us to decipher and apply the MLLE3^Rrm4^ binding code.

## Results

### MLLE3Rrm4 constitutes a novel type of seven-helix MLLE domain

In our previous study, we determined a structural model of MLLE3^Rrm4^ using TopModel, revealing the prototypical five α-helical architecture (helix α1 to α5, Fig. 1B; 14). However, crystallization trials to obtain experimental structural insights by X-ray crystallography failed. Interestingly, structure prediction using AlphaFold (29, 30) suggested the potential presence of two additional helices at the N-terminus of the MLLE3 domain connected via a serine rich flexible linker region (αI and αII N-terminal of α1; Fig. 1B, EV1F).

Intrigued by this, we expressed this N-terminally extended version of MLLE3^Rrm4^ (aa position 679-792) in *E. coli* and purified it to homogeneity (Fig. 1A, Fig. EV1A-B, H-Rrm4-M3 carrying an N-terminal hexa-histidine tag; Materials and methods). Crystallization of the protein was unsuccessful, but upon addition of synthetic peptides of PAM2L1^Upa1^ (aa position 237-254) or PAM2L2^Upa1^ (aa position 944-961; Fig. 1A), crystals of suitable quality for data collection were obtained. The crystals diffracted to 1.7 Å for MLLE3^Rrm4^-PAM2L1^Upa1^ and 2.4 Å for MLLE3^Rrm4^-PAM2L2^Upa1^ (Supporting information (SI) Table S1) and could be solved by molecular replacement using the previously mentioned structural model obtained by AlphaFold. The X-ray structure of MLLE3^Rrm4^ is remarkably similar to the AlphaFold predicted model (RMSD 0.5 Å), confirming the presence of two additional helices (αI and αII) N-terminal to the canonical MLLE domain fold consisting of five α-helices (α1 – α5; Fig. 1B).

To test whether the additional N-terminal αI and αII helices play a role in the recognition of the PAM2-like sequences, we performed GST pull-down assays (14; Materials and methods). To this end, MLLE3^Rrm4^ versions were expressed as a fusion protein with an N-terminal GST tag (Fig. 1A; G-Rrm4-M3-5H, G-Rrm4-M3-7H). G-Pab1-M containing the MLLE domain of Pab1 served as a control. PAM2 and PAM2L motifs of Upa1 were expressed as a fusion protein with the N-terminal hexa-histidine-SUMO tag (Fig. 1A; HS, 17-18 amino acid long peptides, Materials and methods).

GST pull-down experiments revealed that the seven-helix version was able to bind both PAM2L1,2 sequences of Upa1 in a specific manner (Fig. 1C; lane 4, Fig. EV1C, lane 4). In contrast, the shorter five-helix version failed to recognize the PAM2L1,2 sequences (Fig. 1C; lane 3). As expected, MLLE^Pab1^ did not bind the PAM2L1,2^Upa1^ but recognized its cognate PAM2 sequence of Upa1 (Fig. 1C; lane 2; Fig. EV1C, lane 2).

To gain insights into the interaction kinetics, we used ITC using purified proteins to investigate the thermodynamics of MLLE3^Rrm4^ interactions with PAM2L1,2^Upa1^ peptides (Fig. 1A, H-Rrm4-M3; Fig. EV1A). This revealed a K_D_ of 15.7 μM for PAM2L1^Upa1^ and 5.7 μM for PAM2L2^Upa1^ and no binding to PAM2^Upa1^ (Fig. 1D, Fig. EV1D). Thus, our *in vitro* binding results confirmed that the two newly identified N-terminal helices are essential for the interaction and that MLLE3^Rrm4^ consisting of 7 helices, is sufficient for PAM2L1,2^Upa1^ binding with an affinity comparable to the previously tested longer version of Rrm4 (14).

Sequence comparison and AlphaFold predictions with Rrm4 orthologs from related fungi revealed that the seven-helix version of MLLE3^Rrm4^ is conserved in basidiomycetes (Fig. EV1E-F). In distantly related fungi such as the arbuscular mycorrhizal fungus *Rhizophagus irregularis*, we predominantly find the five helix type MLLE domains, suggesting that this short version represents the ancient form (Fig. EV1E-F). In summary, our findings reveal that MLLE3^Rrm4^ possesses two extra N-terminal structural helices crucial for *in vitro* ligand recognition, providing insights into a new type of MLLE domain featuring seven instead of five helices.

### MLLE3^Rrm4^ recognizes PAM2-like sequences with a defined binding pocket

To gain insights into the MLLE3^Rrm4^ – PAM2L1,2^Upa1^ binding mechanism, we inspected our structural model in more detail. Superposition of both complex structures revealed striking similarity of the bound peptide conformations (Fig. 2A, RMSD 0.4 Å). The complex interface features an extensive network of hydrophobic and polar interactions. In both structures, PAM2L^Upa1^ peptides were bound non-canonically to the hydrophobic groove between helices α2 and α3 of the MLLE3^Rrm4^ domain by inserting the bulky aromatic side chains of phenylalanine and tyrosine (Fig. 2A, F248 and Y250 in PAM2L1^Upa1^, F955 and Y957 in PAM2L2^Upa1^). Electron density was present only for the C-terminal half of both PAM2L sequences consisting of 9 or 10 aa residues of the PAM2L1 or PAM2L2 peptides, respectively (Fig. 2A; aa positions 246-254 or 953-962). The N-terminal part of both peptides could not be resolved in the electron density and appears to be flexible.

**Figure 2.**
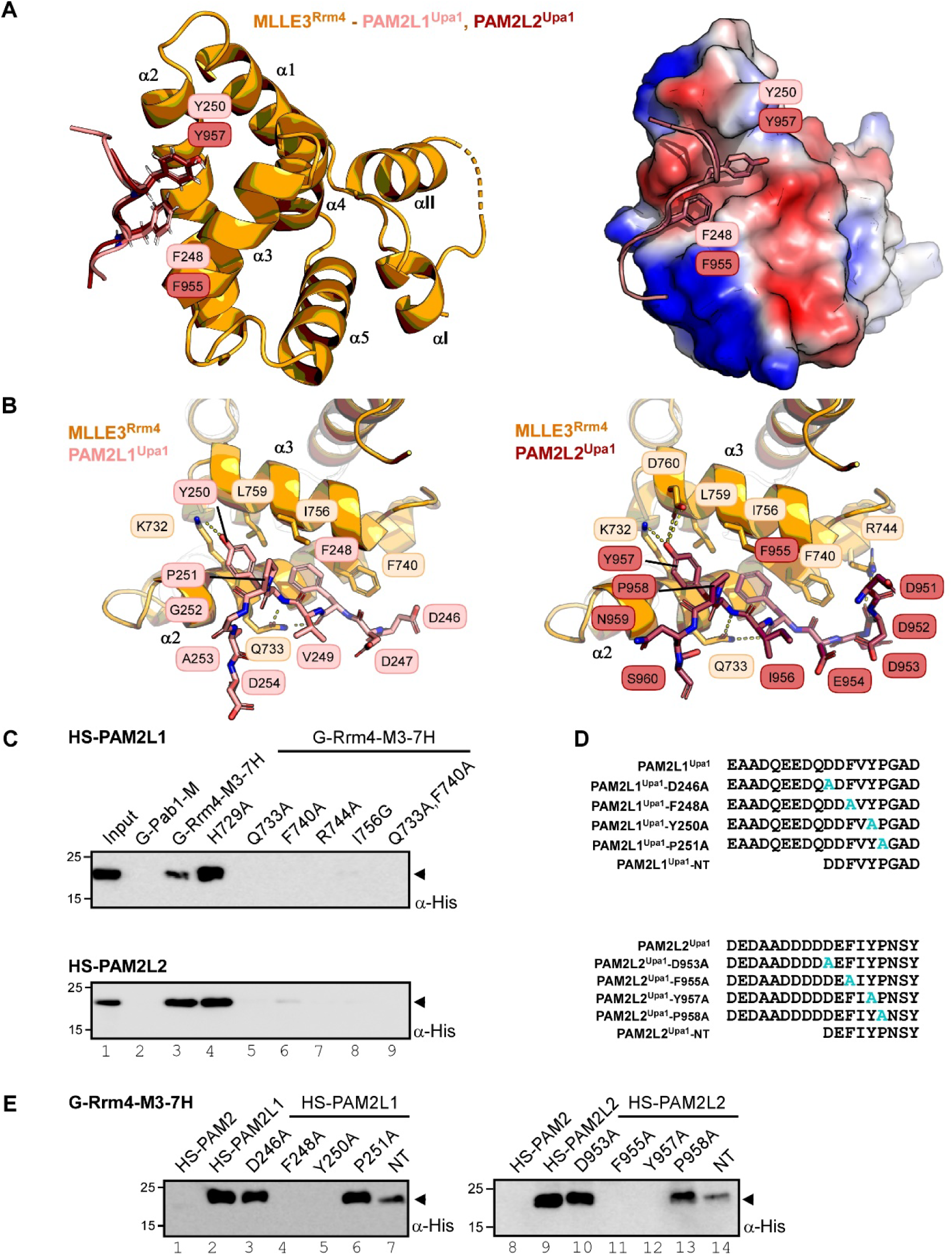
PAM2L ligands are recognized via a new interaction interphase of MLLE3. (**A**) Crystal structures of MLLE3^Rrm4^-PAM2L1^Upa1^, MLLE3^Rrm4^-PAM2L2^Upa1^ complexes are superimposed (RMSD 0.4 Å). The PAM2L1,2^Upa1^ peptides are inserted into the hydrophobic pocket formed by the helices α2, 3 of MLLE3^Rrm4^. Models are represented as a cartoon (left, orange, MLLE3^Rrm4^) and surface (right, MLLE3^Rrm4^, according to electrostatic potential: blue, positively charged; red, negatively charged residues), salmon sticks, PAM2L1^Upa1^; ruby red sticks, PAM2L2^Upa1^. Key residues are labeled. (**B**) Interface between the MLLE3^Rrm4^ and PAM2L1^Upa1^ (left) MLLE3^Rrm4^ and PAM2L2^Upa1^ (right). PAM2L1,2^Upa1^ peptides and interacting side chains of MLLE3^Rrm4^ are shown as sticks. Dashed lines indicate hydrogen bond interactions. (**C, E**) Western blot analysis of GST pull-down experiments using α-His for detection (input, respective His_6_-SUMO peptides). (**D**) Sequence of PAM2L1^Upa1^ and PAM2L2^Upa1^ peptide versions tested in E. Alanine substitution is denoted in cyan.

Hydrophobic residues (G736, F740, P752, I756, and L759) of helices α2-α3 form the core of the peptide binding pocket (Fig. 2A-B). Q733 acts as the coordinator residue, stabilizing the PAM2L^Upa1^ interaction by forming hydrogen bonds with the peptide backbones of the two key bulky aromatic residues F248 and Y250 or F955 and Y957 in the case of PAM2L1^Upa1^ or PAM2L2^Upa1^, respectively (Fig. 2A-B). Furthermore, the positively charged side chain of K732 in α2 of MLLE3^Rrm4^ establishes a polar contact with the hydroxyl group of Y250 of PAM2L1^Upa1^ (Fig. 2B, PAM2L1^Upa1^), whereas the negatively charged side chain of D760 in α3 contacts the hydroxyl group of the Y957 of PAM2L2^Upa1^ (Fig. 2B, PAM2L2^Upa1^).

For verification, we altered key residues of MLLE3^Rrm4^ and tested the resulting constructs in GST pull-down assays (Fig. 2C; Materials and methods). Variations at positions Q733A, F740A, R744A and I756G in MLLE3^Rrm4^ strongly affected the interaction with both HS-PAM2L1,2^Upa1^ versions (Fig. 2C, lane 5-8). The double substitution Q733A,F740A showed no binding (Fig. 2C, lane 9). In contrast, substituting H729A did not alter the interaction indicating that this position is not essential for interaction (Fig. 2C, lane 4).

### Identifying an essential FxY core in PAM2-like peptides of Upa1

To determine the critical residues in the PAM2L1,2^Upa1^ peptides, different variants were tested in GST pull-down assays (Fig. 2D). Alanine substitutions at positions F248 or Y250 in HS-PAM2L1^Upa1^ as well as F955 or Y957 in HS-PAM2L2^Upa1^ abolished the interaction with the G-Rrm4-M3-7H (Fig. 2E, lane 4, 5 and lane 11, 12, respectively). This outcome was expected since F248, Y250 in PAM2L1 and F955, Y957 in PAM2L2^Upa1^ are inserted into the hydrophobic pocket between helices 2 and 3. F248 of PAM2L1^Upa1^ and F955 of PAM2L2 ^Upa1^ are involved in aromatic stacking with F740 in the MLLE3^Rrm4^ similar to the PAM2^PAIP2^-MLLE^PABP^ interaction in human, which is crucial for peptide binding (22). Alanine substitutions at positions P251 in PAM2L1^Upa1^ (P958 in PAM2L2^Upa1^) did not affect the binding with the MLLE3^Rrm4^ (Fig. 2D, lane 6, 13). This is consistent with the structural information, since the proline residues of both PAM2L1,2^Upa1^ are exposed outside the peptide binding pocket and do not significantly contribute to PAM2L^Upa1^-MLLE3^Rrm4^ interactions (Fig. 2A-B). Binding was not affected when D246 in PAM2L1^Upa1^ and D953 in PAM2L2^Upa1^ were substituted with alanine (Fig. 2E, lane 3, 10). High b-factors reveal a high degree of flexibility in this region of both PAM2L1,2^Upa1^ peptides (Fig. EV2E). Therefore, D246 and D953 might not be essential for interaction but could play a supporting role in stabilizing the complex through electrostatic interactions with R744 of MLLE3^Rrm4^ (Fig. 2B). Additionally, GST pull-down assays using an N-terminally truncated (NT) versions of both HS-PAM2L1 and 2^Upa1^ confirmed that the shorter version of the peptides found in the co-crystallized structure (Fig. 2A) is sufficient for the interaction (Fig. 2E, lane 7, 14).

Previously, we reported that the Rrm4 from *Rhizophagus irregularis* (*Ri*Rrm4) co-localized with Upa1-Gfp and shuttled on endosomes in *U. maydis* (31). We hypothesized that this interaction might be mediated by the MLLE3 domain of *Ri*Rrm4 (*Ri*MLLE3^Rrm4^) by recognising the PAM2L motifs of Upa1. However, contrary to the MLLE3^Rrm4^ of *U. maydis* (*Um*MLLE3^Rrm4^), *Ri*MLLE3^Rrm4^ consists of only five helices. Comparing the AlphaFold structural model of the complex of *R. irregularis* MLLE3^Rrm4^ with *U. maydis* PAM2L2^Upa1^ peptide (ipTM = 0.7) revealed that *Ri*MLLE3^Rrm4^ retains a perfectly similar PAM2-like peptide binding pocket comparable to *Um*MLLE3^Rrm4^ in spite of having only five helices (Fig. EV2C). Additionally, we observed that the αI and αII of *Um*MLLE3^Rrm4^ are involved in multiple intra molecular interaction and pushing the α5 towards the α3, whereas in *Ri*MLLE3^Rrm4^ the α5 is slightly deviated and aligned closely towards the α3 even in the absence of the αI and αII (Fig. EV2 D). To verify this interaction, we performed pull-down experiments. Consistently, HS-PAM2L1,2^Upa1^ peptides from *U. maydis* bound to the five-helix version of *Ri*MLLE3^Rrm4^ (Fig. EV2B, G-*Ri*Rrm4-M3, lane 2, 3). Hence, the MLLE3-PAM2L interaction appears to be evolutionarily conserved. In essence, structural and biochemical analysis allowed the identification of the FxY core of PAM2L sequences as essential determinant for MLLE3^Rrm4^ recognition.

### MLLE3^Rrm4^ is necessary and sufficient for endosomal attachment

For functional analysis of the seven-helix type MLLE3 domain, we expressed various versions of Rrm4 (Fig. 3A), fused with the red fluorescent protein mKate2 at their C-termini in *U. maydis* (designated Kat, 32). As a genetic background, we used laboratory strain AB33 expressing bE/bW, the key heteromeric transcription factor for hyphal growth, under the control of the nitrogen source-regulated promoter P_nar1_ (33). Therefore, unipolar hyphal growth can be elicited efficiently and synchronously by switching the nitrogen source (11, 34). Additionally, the strains expressed Upa1-Gfp to verify that, as expected, mutations in Rrm4 did not influence shuttling of Upa1-positive endosomes (Fig. EV3A; 14, 15).

**Figure 3.**
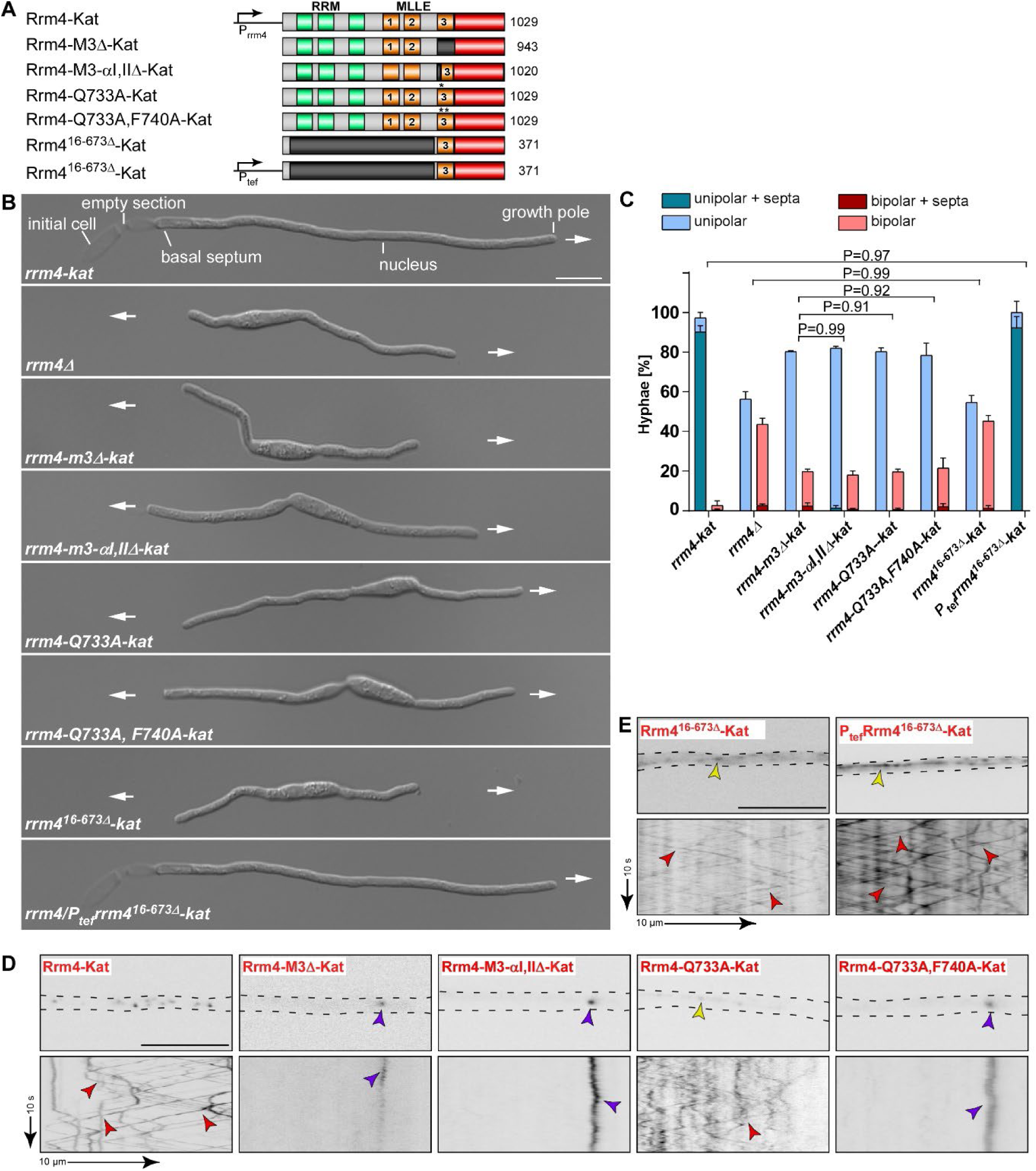
A seven-helix type MLLE domain is necessary and sufficient for endosomal attachment. **(A**) Schematic representation of Rrm4 variants (amino acid number indicated; drawn not in scale): dark green, RNA recognition motif (RRM); orange, MLLE domains; red, mKate2. (**B**) Hyphal growth of AB33 derivatives (6 h.p.i.; size bar 10 μm). Growth direction is indicated by arrows. (**C**) Quantification of hyphal growth of AB33 derivatives shown in panel B (6 h.p.i.): unipolarity, bipolarity and basal septum formation were quantified (error bars, SEM.; n = 3 independent experiments, > 150 hyphae were counted per strain; for statistical evaluation, the percentage of uni– and bipolarity was investigated and unpaired two-tailed Student’s t-test was performed (α<0.05). (**D-E**) Micrographs (inverted fluorescence image; size bar, 10 μm) and corresponding kymographs of AB33 hyphae derivatives (6 h.p.i.) showing movement of Rrm4-Kat variants in hyphae (inverted fluorescence images; arrow length on the left and bottom indicates time and distance, respectively). Processive signals, aberrant microtubule staining and accumulation of static Rrm4-Kat signals are indicated by red, yellow and purple arrowheads, respectively.

The wild type version of Rrm4-Kat is fully functional, as indicated by unipolar growth and shuttling on endosomes (Fig. 3B-D). As reported previously, deletion of MLLE3^Rrm4^ (Rrm4-M3Δ-Kat; 35) resulted in bipolar growth and static Rrm4 signals (Fig. 3A-D; Fig. EV3B-D; 14). Rrm4 versions lacking αI and αII also lost functionality (Fig. 3A-D; Fig. EV3B-D). Similarly, double substitutions in key amino acids Q733A and F740A affected function (Fig. 3A-D; Fig. EV3B-D). Interestingly, the variation of Q733 to alanine caused the characteristic increase of bipolar growth indicating that the protein is not functional (Fig. 3B-C). However, weak shuttling of Rrm4-Q733A-Kat was observed, suggesting residual interaction with PAM2L motifs of Upa1 (Fig. 3D; Fig. EV3A-C). Taken together, *in vivo* studies are in agreement with our structural data and highlight the importance of MLLE3^Rrm4^ for endosomal transport.

To demonstrate that MLLE3^Rrm4^ alone is sufficient for endosomal shuttling, we generated a strain expressing only MLLE3^Rrm4^ (Fig. 3A). We tested two versions. Firstly, in Rrm4^16-673Δ^-Kat strains, MLLE3^Rrm4^ was expressed under the control of the native promoter. Hyphae grow bipolarly because this domain cannot replace the full-length protein in function (Fig. 3B-C). Secondly, the MLLE3^Rrm4^ construct used above was expressed under the control of the constitutively active tef promoter by ectopic insertion at the i*p^S^* locus, in addition to the wild type allele of *rrm4* (Fig. 3A, see Materials and methods). As expected, this strain showed unipolar growth (Fig. 3B,C). Importantly, in both strains, the MLLE3^Rrm4^-Kat fusion exhibited endosomal shuttling with comparable velocity and distance travelled similar to the wild type (Fig. 3E, Fig. EV3D-F). In both cases, MLLE3-Kat was mislocalized and exhibited aberrant staining of microtubules (Fig 3E). This is reminiscent for Rrm4 versions lacking the accessory function of MLLE1 and MLLE2 (14). In essence, the seven-helix type MLLE3 domain of Rrm4 is necessary and sufficient for attachment to transport endosomes.

### Deciphering the binding code for MLLE^Pab1^ and MLLE3^Rrm4^

To understand how MLLE^Pab1^ and MLLE3^Rrm4^ domains of *U. maydis* recognize the PAM2 and PAM2L sequences of Upa1, we begun by solving the co-structure of MLLE^Pab1^ with the cognate ligand PAM2^Upa1^. Initially, we employed AlphaFold to predict the structural model of MLLE^Pab1^, revealing that it is consisted of five helices similar to the MLLE^PABPC1^ (Fig.EV4D, 16). This structural similarity is reflected in sequence similarity to other MLLE domains (Fig. EV4E), indicating evolutionary conservation from fungi to mammals.

For experimental verification, we expressed a version of MLLE^Pab1^ (aa position 567-636) in *E. coli* and purified to homogeneity (Fig. EV4A-B; H-Pab1-M carrying an N-terminal hexa-histidine-tag; Materials and methods). To gain insights into the interaction kinetics, we utilized ITC with purified protein to investigate the thermodynamics of MLLE^Pab1^ interaction with the PAM2^Upa1^ peptide (aa position 128-144; Fig. 1A). We determined a K_D_ of 15 μM (Fig. EV4C), confirming that this version is sufficient for PAM2 binding with an affinity comparable to our previous report (14).

Once again, attempts to crystallize the protein in its apo state were unsuccessful. However, upon addition of synthetic PAM2^Upa1^ peptide, crystals of suitable quality for data collection were obtained. These crystals diffracted to a resolution of 2.0 Å and were successfully solved by molecular replacement using the structural model generated by AlphaFold (Fig. EV4D; SI Table S1, MLLE^Pab1^-PAM2^Upa1^).

The peptide bound to MLLE^Pab1^ by wrapping around α3 of MLLE^Pab1^,interacting with the hydrophobic pockets between helices α2-α3 and helices α3-α5 (Fig. 4A). Hydrophobic residues in the α2, α3 and α5 of MLLE^Pab1^ form the core of its peptide binding pocket (Fig. 4A). These two pockets provide the most important binding interactions, recognizing the conserved leucine (L132) and phenylalanine (F139) residues in the PAM2^Upa1^ peptide (Fig. 4A). Notably, the conserved Y580 within MLLE^Pab1^ restricts the second hydrophobic pocket, thereby encompassing F139 (Fig. 4A). Several hydrogen bonds stabilize the interaction: e.g., N135 and A136 of PAM2^Upa1^ interact with K593 of MLLE^Pab1^ and S142 of PAM2^Upa1^ interacts with E577 of MLLE^Pab1^. Additionally, P141 establishes a hydrogen bond to the coordinator glutamine (Q573) of MLLE^Pab1^ (Fig. 4 A).

**Figure 4.**
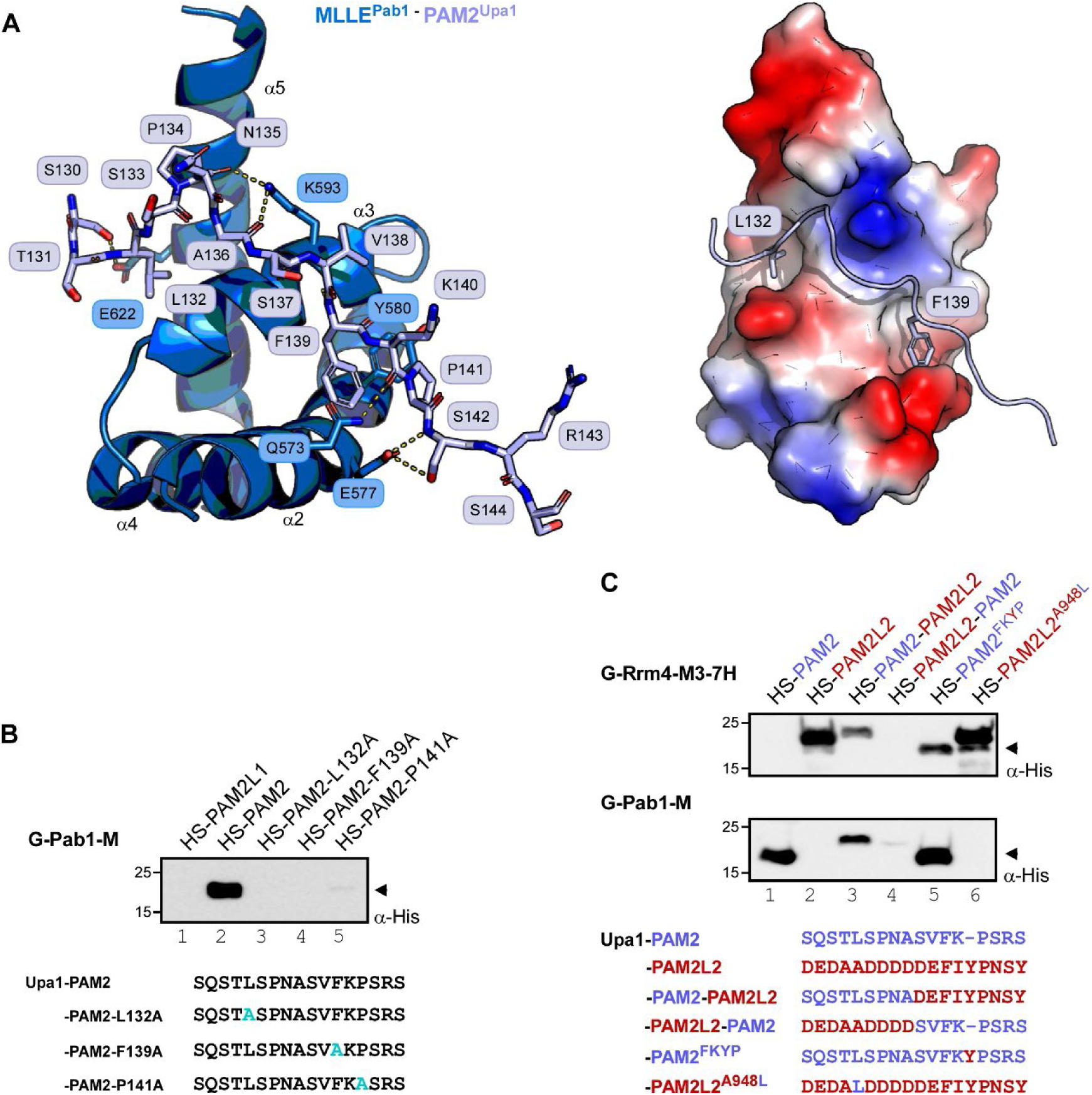
The MLLE domain of Pab1 recognizes its PAM2 ligand in a canonical fashion. (**A**) Crystal structure of PAM2^Upa1^ bound to MLLE^Pab1^. The PAM2^Upa1^ peptide wraps around the MLLE^Pab1^. Key residues (L132 and F139) are inserted into the hydrophobic pocket formed between the helices α3,5 and between the helices α2,3. Models are represented as a cartoon (left) and surface (right). Light blue sticks, PAM2^Upa1^ peptide; interacting side chains of MLLE^Pab1^ are shown as sticks and dashed yellow lines indicate hydrogen bonding. (**B-C**) Western blot analyses of GST pull-down experiments using α-His for detection. PAM2^Upa1^ sequence is denoted in light blue, PAM2L2^Upa1^ sequence is denoted in red, hybrid versions are denoted in light blue-red dual color)

For validation, we introduced variations in the key residues of the PAM2^Upa1^ peptide and tested the resulting constructs in GST pull-down assays (Fig. 4B, Materials and methods). Variations at positions L132A or F139A in HS-PAM2^Upa1^ abolished the interaction with the G-Pab1-M (Fig. 4B, lane 3, 4), as expected, since these residues are crucial for the MLLE binding (Fig. 4A, Fig. EV4E). Substitution at P141A disrupted the interaction with G-Pab1-M (Fig. 4B, lane 5). This is supported by the b-factors, which are low for P141 (Fig. EV4F), indicating low flexibility that might increase upon variation to alanine. Our *in vitro* interaction assays confirmed that L132 and F139 are the crucial residues in PAM2^Upa1^ for interaction with MLLE^Pab1^. Thus, we observed a clear difference in the mode of binding between PAM2^Upa1^ and PAM2L^Upa1^ by the MLLE domains of Pab1 and Rrm4, respectively.

To further confirm the binding code of the two types of MLLE domains, we generated hybrid versions of the PAM2^Upa1^ and PAM2L^Upa1^ and tested the resulting constructs in GST pull-down experiments. The hybrid containing the N-terminal half of PAM2^Upa1^ with the critical leucine and the C-terminal tyrosine was bound by both MLLE domains, but showed reduced signal intensity (Fig. 4C, lane 3). In contrast, the complementary hybrid sequence was not bound by any MLLE domain (Fig. 4C, lane 4). Even more convincingly, the simple insertion of the critical tyrosine of the PAM2L^Upa1^ sequence in the PAM2^Upa1^ background resulted in a synthetic version that is recognized by both MLLE domains (Fig. 4C, lane 5), whereas insertion of critical leucine residue in the PAM2L^Upa1^ peptide did not mediate the peptide interaction with MLLE^Pab1^ (Fig. 4C, lane 6). In essence, solving the co-structures of MLLE domains with their cognate PAM2 and PAM2L sequences allowed deciphering binding specificity and revealed two distinct, evolutionarily conserved peptide recognition modes.

### Identification of new Rrm4 interaction partners

As pointed out above, Rrm4 remains associated with shuttling endosomes in the absence of Upa1, suggesting the presence of additional endosomal adaptor proteins (15). Therefore, we aimed to leverage our understanding of the binding mechanism and critical residues of PAM2L recognition to predict unknown interaction partners of MLLE3^Rrm4^. A similar strategy, based on the PAM2 consensus sequences, previously identified Upa1 and Upa2 as Rrm4 interaction partners (15, 34).

Initially, we utilized the fact that the critical FxY core sequence of PAM2L1,2^Upa1^ is N-terminally flanked by acidic residues (Fig. 1A). Using the KEGG motif search algorithm (36), we screened for similar PAM2L-motif containing proteins in *U. maydis* genome using the code ([DE]-[DE]-F-x-Y), retrieving 47 candidates, including the known PAM2L1,2^Upa1^. Next, we assessed the accessibility of the interaction motif by visually scanning AlphaFold-predicted models in the unstructured regions (16, 37). We shortlisted 23 candidates containing PAM2L motifs in either intrinsically disordered regions (IDR) or short unstructured linkers. Third, we examined the evolutionary conservation of the PAM2L sequences in fungi and shortlisted 12 candidates (SI, Table S4). Finally, we searched for candidates in which the PAM2L motif is present in basidiomycetes with hyphal growth mode, such as *U. hordei* and *Sporisorium reilianum,* but absent in those proliferating mainly in the yeast form, such as *Malassezia globosa* and *Cryptococcus neoformans*. Applying these four levels of selection criteria, we identified nine new PAM2L candidates specific for basidiomycetes forming hyphae. The remaining three candidates harbored PAM2L sequences in linker regions (SI, Table S4).

Initially, we selected Vps8 (Fig. 5A; UMAG_15064), which is part of the CORVET complex (class C core vacuole/endosome tethering) and is known to be present on Rab5a positive endosomes in *U. maydis* (38). Examining the evolutionary conservation of the PAM2L sequence revealed a potential second PAM2L sequence, containing a consensus FxY core motif (Fig. EV5A). Both sequences are present in its IDR (Fig. 5A,C; PAM2L1^Vps8^, PAM2L2^Vps8^) reminiscent of Upa1 which features two PAM2L sequences in its N-terminal unstructured region (Fig. EV5C, 15), absent in the respective *M. globosa* homologs (Fig. EV5A).

**Figure 5.**
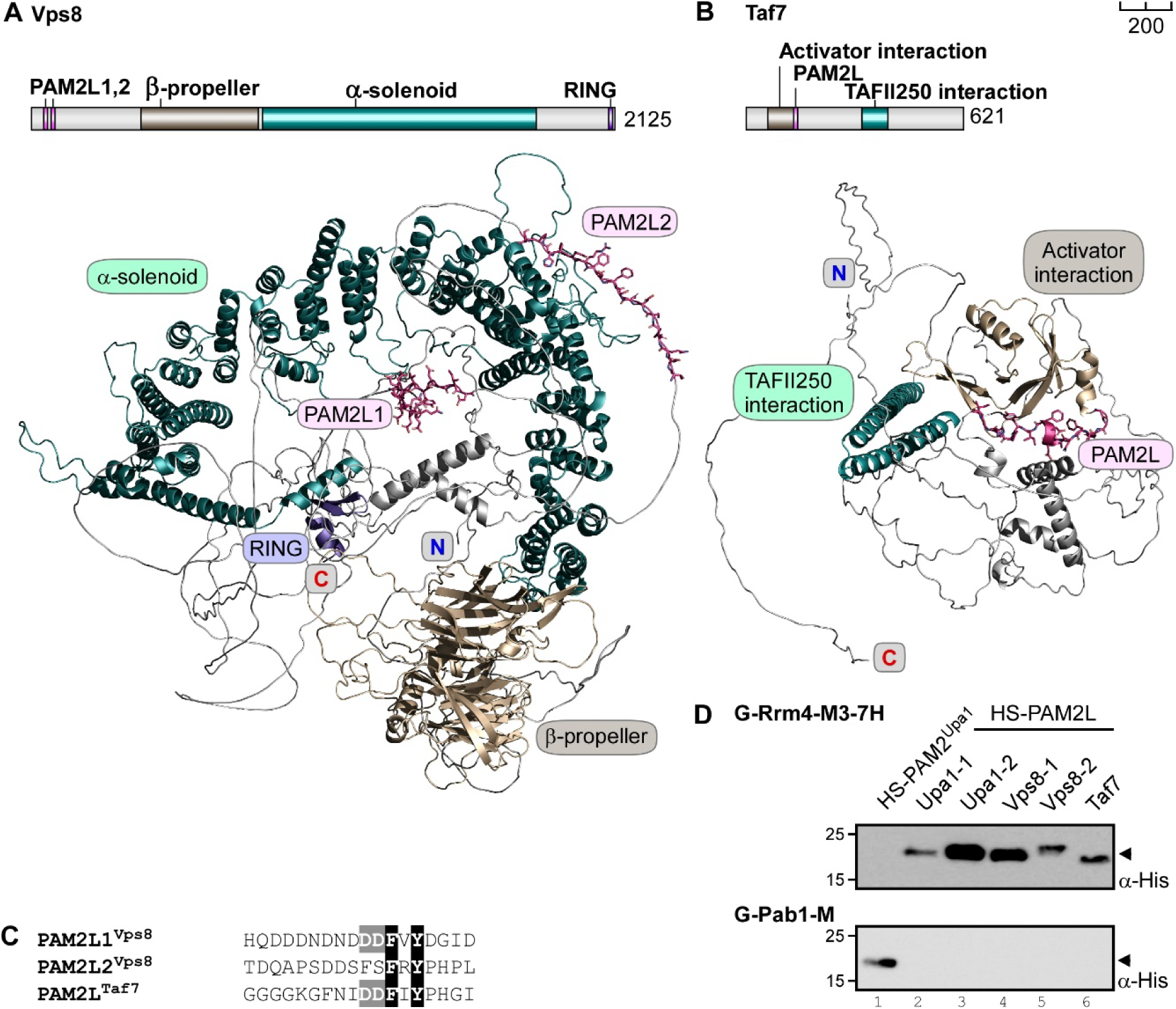
Vps8 and Taf7 are *de novo* predicted interaction partners of Rrm4. (**A**) Schematic representation of Vps8 (UMAG_15064) (top) (aa number is indicated next to protein bars, drawn to scale, bar on top right is 200 AA; pink, PAM2L1,2; wheat, β-propeller domain; dark teal, α-solenoid domain; purple, RING domain. 3D structural model predicted using AlphaFold (bottom) with domains depicted using the above color code, amino (N), carboxy (C) terminals are indicated. (**B**) Schematic representation of Taf7 (UMAG_10620) drawn to scale (Pink, PAM2L; wheat, activator interaction domain; dark teal, homolog of human TAFII250 interaction domain). Structural model predicted using AlphaFold (bottom) with domains depicted using the above color code, amino (N), carboxy (C) terminals are indicated. (**C**) De novo predicted PAM2L peptides of Vps8 and Taf7 are denoted, conserved, crucial resides are shaded in black, conserved key acidic residues are shaded in grey (**D**) Western blot analyses of GST pull-down experiments using α-His for detection.

To test a PAM2L candidate present in a linker region, we selected Taf7 (Transcription initiation factor TFIID subunit 7, UMAG_10620; Fig 5B, Fig. EV5B). The evolutionarily conserved Taf7 functions during transcription and nucleocytoplasmic mRNA export (39, 40).

To evaluate the binding capacity of the *de novo* predicted novel PAM2L-motifs, we performed GST pull-down assays (Fig. 5D; HS-PAM2L1^Vps8^, HS-PAM2L2^Vps8^ and HS-PAM2L^Taf7^, Materials and methods). PAM2L sequences of Vps8 and Taf7 interacted specifically with MLLE3^Rrm4^ but not with MLLE^Pab1^ (Fig. 5C-D; lane 4-6). The second PAM2L sequence of Vps8, lacking acidic residues N-terminal of the FxY core, showed a weaker interaction with MLLE3^Rrm4^ (Fig. 5C-D, lane 5), emphasizing the supportive role of these acidic resides through electrostatic interaction. Thus, we successfully predicted additional interaction partners of MLLE3^Rrm4^ in *U. maydis* and demonstrated their interaction *in vitro*. In essence, we succeeded in deciphering and applying the MLLE binding code to identify novel interaction partners *de novo*. Future research will clarify the mechanistic and functional details of the underlying interactions.

### Human MLLE domains differentiate between binding partners

To explore whether a similar MLLE binding code governs beyond the fungal lineage, we turned our focus to the two known MLLE domains from humans, MLLE^PABPC1^ and MLLE^UBR5^ (Fig. 6A). GST pull-down assays demonstrated that MLLE^PABPC1^ interacts with known PAM2 sequences from PAIP2, TOB and GW182 (Fig 6C-D, lane 1-3). However, we couldn’t detect the known interaction with the PAM2 sequence of MKRN1 (RNA-binding E3 ubiquitin ligase Makorin Ring Finger Protein 1; aa position 161 – 177; 41). Given that a short PAM2^MKRN1^ version of 18 amino acids was not previously tested, additional flanking sequences may be necessary for this interaction. Intriguingly, our analysis of MKRN1 uncovered a potential PAM2 variant (PAM2L^MKRN1^) sequence at position 329-346 that is evolutionarily conserved in a low complexity switch region (Fig. 6 A-C, EV6, PAM2L^Mkrn1^). Indeed, this PAM2L sequence exhibited a weak interaction with MLLE^PABPC1^ (Fig. 6C, lane 5). Consistent with previous reports, MLLE^PABPC1^ did not bind to the PAM2L sequence of UBR5 (Fig. 6D, lane 6; 26).

**Figure 6.**
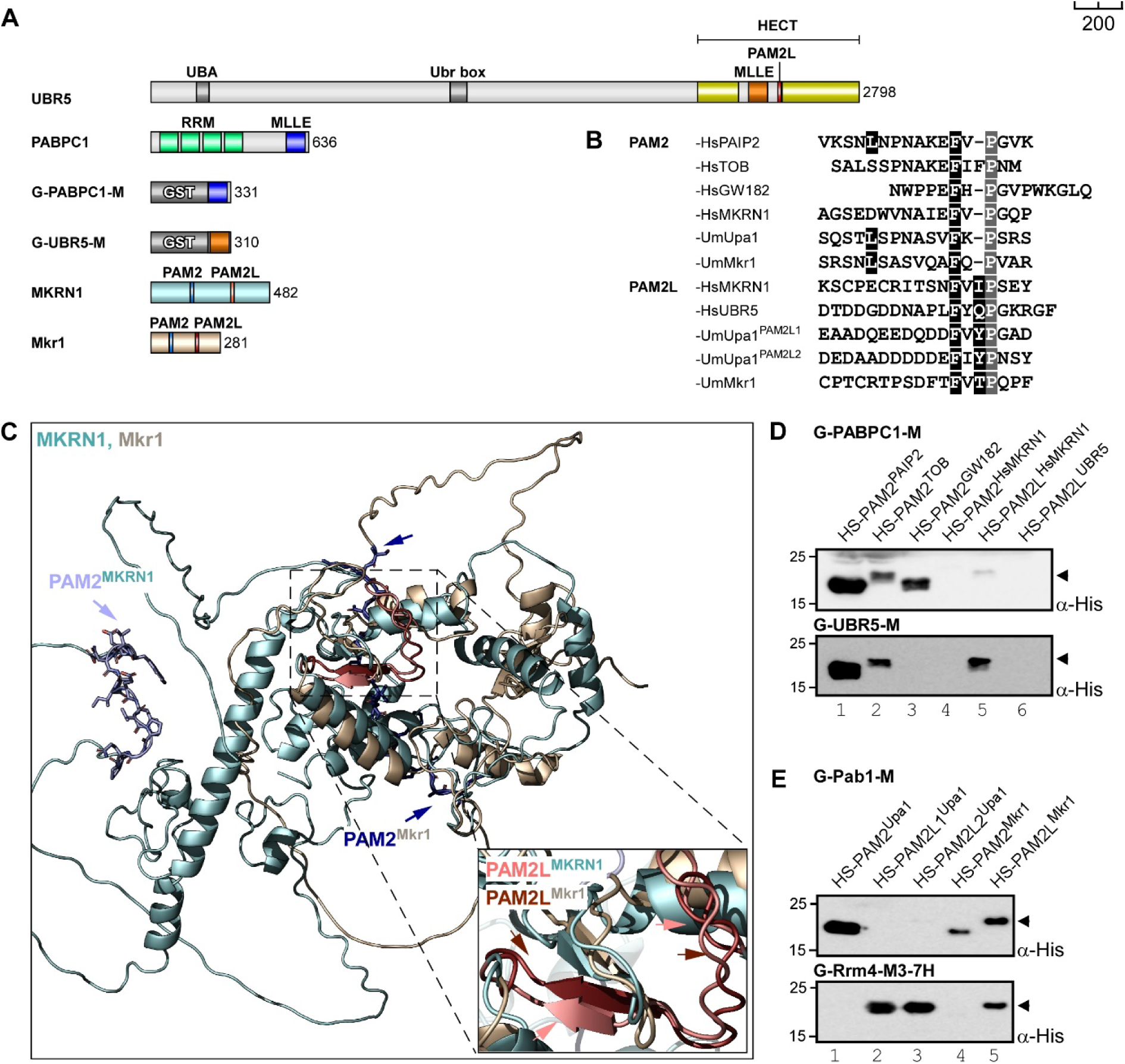
MLLE domains of PABC1 and UBR5 differentiate between binding partners. (**A**) Schematic representation of protein variants drawn to scale (aa number is indicated next to protein bars, drawn to scale, bar on top right is 200 AA; grey, UBA, Ubr box, GST; yellow, HECT; orange, MLLE^UBR5^; red, PAM2L; blue, MLLE^PABC1^; green, RRM; cyan, MKRN1; wheat, Mkr1.(**B**). Comparison of PAM2 and PAM2L sequences found in Upa1 (UniProtKB ID: A0A0D1E015) with those of human proteins, such as PAIP2 (Q9BPZ3), TOB (P50616), GW182 (Q9HCJ0), MKRN1 (Q9UHC7), Mkr1 (A0A0D1E4Z6), UBR5 (O95071). (**C)** Structural models of MKRN1 and Mkr1 predicted using AlphaFold, domains depicted in the color code similar to the respective labels, PAM2L motifs are shown in. (**D-E**) Western blot analysis of GST pull-down experiments using α-His for detection.

Comparative analysis with MLLE^UBR5^ showed clear differences in binding specificity. While it recognized classical PAM2 sequences of PAIP2 and TOB, it did not interact with the non-canonical PAM2^GW182^ variant (Fig. 6D, lane 1-3; G-Ubr5-M). Furthermore, MLLE^UBR5^ exhibited stronger binding to the PAM2L^MKRN1^ than MLLE^PABPC1^ (Fig. 6D, lane 5; G-Ubr5-M). Although we failed to detect the interaction with its own PAM2L^HECT^ sequence (Fig. 6D, lane 6; G-Ubr5-M), this failure is likely attributable to its low binding affinity (*K*_D_ of 50 µM, 26). Therefore, human MLLE domains display differential binding capacities, likely utilizing specific binding interfaces. Notably, we identified MKRN1 as a new interaction partner of UBR5 (see Discussion).

Given the significance of RNA-binding MKRN1 as a target for MLLE domain-containing proteins in human, we investigated the MKRN1 homologue Mkr1 from *U. maydis* (UMAG_12122; Fig. 6C, EV6). Mkr1 contains a conserved PAM2 and a PAM2L sequence at position 77 – 93 and 180 – 197, respectively, in a conserved low complexity switch region akin to its human ortholog (Fig. 6A-C, Fig. EV6). GST pull-down experiments revealed that MLLE3^Rrm4^ only bound the PAM2L^Mkr1^ sequence (Fig. 6E, lane 4, 5; G-Rrm4-M3-7H, lane 4,5), whereas MLLE^Pab1^ recognized both PAM2^Mkr1^ and PAM2L^Mkr1^ sequences (Fig. 6E, G-Pab1-M, lane 4, 5). This confirms that PAM2^Mkr1^ functions as classical PAM2 motif. However, the PAM2L^Mkr1^ sequence is recognized by both MLLE domains, reminiscent of our synthetic PAM2/PAM2L peptides (Fig. 4C, lane 3, 5) and the PAM2L^MKRN1^ binding by human MLLE domains (Fig. 6 D, lane 5). In summary, Mkr1 is a previously unknown interaction partner of the endosomal mRNA transporter Rrm4 as well as the poly(A)-binding protein Pab1. Overall, MLLE domains serve as sophisticated binding platforms for the formation of defined interaction networks with PAM2 and PAM2L motif-containing partners.

## Discussion

The transport of mRNAs via endosomes represents a fundamental trafficking mechanism across various organisms, including fungi, plants and humans (6). Moreover, growing evidence suggests that endosome-coupled translation is a conserved biological process crucial for local protein synthesis serving purposes such as septin complex formation in hyphae and localized mitochondrial protein import in neurons (10, 42). To elucidate the underlying mechanisms, it is essential to comprehend how mRNAs are attached to endosomes. Here, we present that the endosomal mRNA transporter Rrm4 harbors a novel seven-helix-type MLLE domain, which is necessary and sufficient for endosomal attachment (Fig. 7A). These underlying interactions are integral components of a sophisticated MLLE/PAM2 recognition system, forming a resilient SLiM-based network with a high level of binding redundancy (Fig. 7B).

**Figure. 7.**
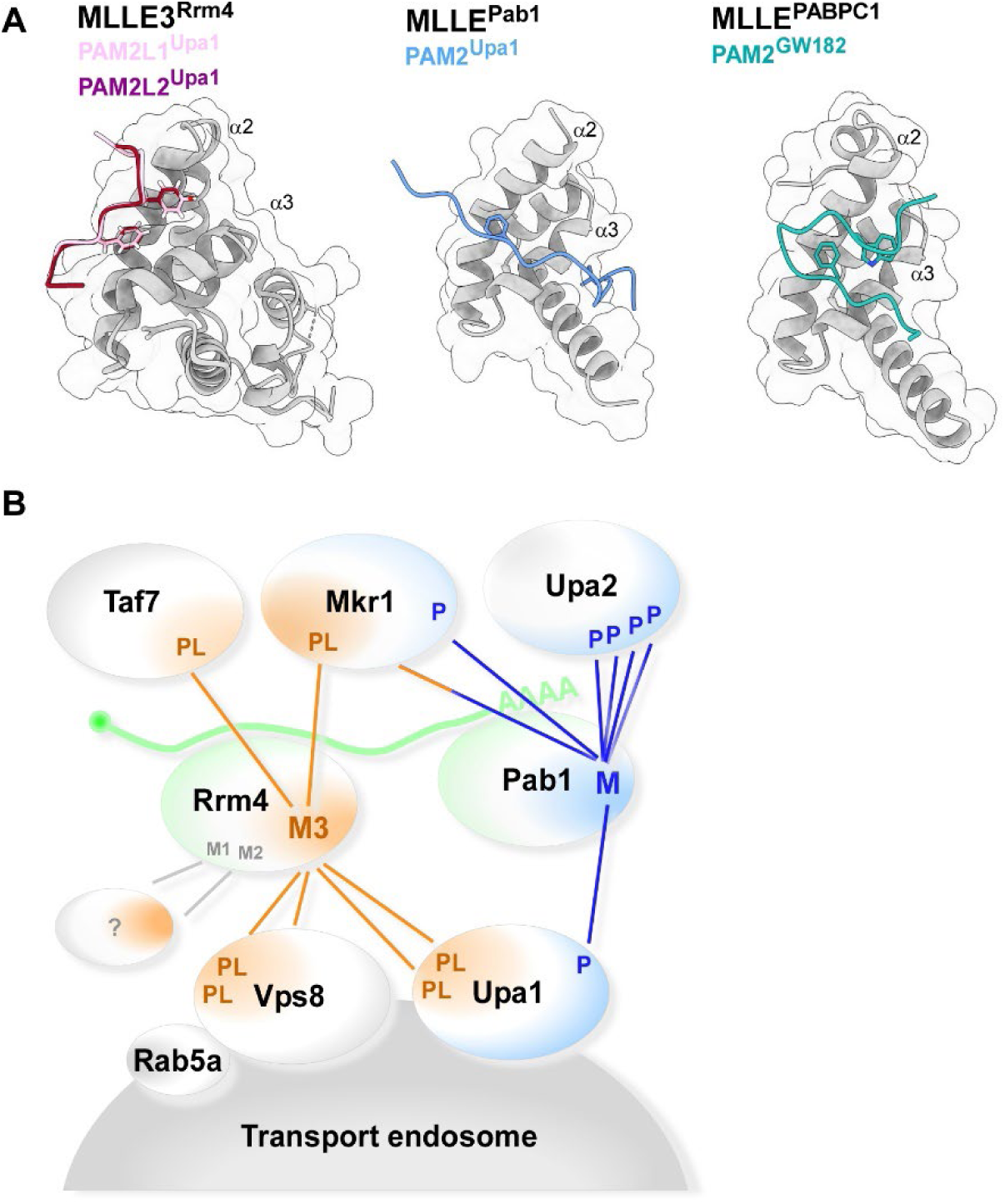
MLLE domains exhibit defined binding specificity. (**A**) Structures of three different MLLE domains bound to ligands. MLLE3^Rrm4^-PAM2L1,2^Upa1^, MLLE^Pab1^ –PAM2^Upa1^ and MLLE^PABPC1^-PAM2^GW182^ (right, PDB ID: 3KTP). Helices 2 and 3 are labeled. (**B**) Model depicting the complex protein-protein interaction network based on the binding specificity of MLLE domains of Rrm4 (orange) and Pab1 (blue). The following symbols are used: M1, MLLE1; M2, MLLE2; M3, MLLE3; M, MLLE; PL, PAM2-like; P, PAM2; mRNA with poly(A) tail in green; ?, unknown proteins.

### The SLiM-based MLLE domain binding code

The canonical MLLE domain is typically comprised of four to five helices arranged in a defined architecture, with a specific binding interface involving helices 2,3 and 5 recognizing cognate PAM2 motifs (Fig. 7A). An exception to this is the PAM2 sequence of GW182, which utilizes a different interface along helix 2-3 (Fig. 7A, right; 23). Here, we elucidate how the new seven-helix-type MLLE domain, MLLE3^Rrm4^ is able to differentiate between PAM2 and related PAM2L sequences. MLLE3^Rrm4^ contains two additional helices at the N-terminus of the conserved MLLE core, a feature absent in all currently known MLLE domains: MLLE1^Rrm4^, MLLE2^Rrm4^, MLLE^Pab1^ from *U. maydis*, and MLLE^PABC1^ and MLLE^UBR5^ domains from human and other eukaryotes (Fig. 7A; 14, 19). The two additional helices αI and αII of MLLE3^Rrm4^ are not directly involved in peptide recognition but rather prevent canonical PAM2 binding. The seven-helix-type MLLE3^Rrm4^ utilizes a different binding interface along helix 2, resembling the PAM2^GW182^ binding of human MLLE^PABC1^ (Fig. 7A). Thus, we demonstrate that the conserved α−helical core of the MLLE domain employs various, yet defined, binding pockets to determine a precise specificity of SLiM binding. Additionally, we decoded the central amino acid motifs within the PAM2 and PAM2L sequences that determine exquisite binding specificity.

Intriguingly, fundamental principles of MLLE interactions are conserved in humans. The MLLE domain-containing proteins PABC1 and UBR5 share targets such as PAIP2 and TOB (19). However, we also observed clear differential recognition, such as MLLE^PABC1^ and MLLE^UBR5^ binding to PAM2^GW182^ and PAM2L^MKRN1^, respectively. Recent cryo-EM structural analyses revealed that UBR5 forms functional dimers and tetramers (27, 28). Interestingly, MLLE^UBR5^ is inserted in the middle of the catalytic HECT domain essential for ubiquitin transfer (Fig. 6A; 24, 27, 28). Thus, recruitment of other factors, such as our newly described interaction with E3 ubiquitin ligase MKRN1 via its PAM2L sequence, might directly influence the ubiquitin chain elongation activity of UBR5. The fact that MKRN1 is an RBP links both MLLE domain proteins, UBR5 and PABPC1, to RNA binding protein networks comparable to their fungal counterparts. Consistently, Mkr1 from *U. maydis* contains a PAM2 and PAM2L sequence and might also contribute to endosomal mRNA transport.

### A SLiM-based RNA-binding protein network for endosomal attachment of mRNAs

Understanding endosomal mRNA transport requires clear elucidation of how mRNAs, associated RNA-binding proteins, and ribosomes are tethered to endosomes. Previous studies have shown that Rrm4 continues to hitchhike on endosomes even in the absence of the PAM2L-containing protein Upa1, indicating the presence of additional adaptor proteins. Rrm4 contains a platform of three MLLE domains that operates with a strict hierarchy: MLLE1^Rrm4^ and MLLE2^Rrm4^ play accessory roles, whereas MLLE3^Rrm4^ is essential for endosomal shuttling (Fig. 7B; 14).

Applying the MLLE-binding code led to the identification of new interactors of MLLE3^Rrm4^, such as Taf7 and Vps8 which contain experimentally verified PAM2L sequences (Fig. 7B). Taf7 is a potential homolog of the general transcription factor Taf7 from *S. pombe* (39, 43). It might be loaded onto pre-mRNAs during transcriptional initiation, and its interaction with Rrm4 could be important during remodeling of mRNPs in the cytoplasm prior to endosomal loading. This function mirrors the mRNPs remodeling role of the nuclear factor Loc1p during *ASH1* mRNA transport in *S. cerevisiae* (44). Additionally, the nuclear history of splicing factor Num1 from *U. maydis*, which interacts with molecular motor Kinesin-1, has been implicated in microtubule-dependent trafficking, such as endosomal mRNA transport (45).

The second example, Vps8 is particularly intriguing due to its known localization to transport endosomes through interaction with Rab5a (38). Vps8 performs an evolutionarily conserved function during endosome maturation by recruiting the CORVET complex (class C core vacuole/endosome tethering; 38, 46). In *U. maydis,* Vps8 might have acquired new functionality during endosomal mRNA transport, potentially serving as an additional endosomal adaptor alongside Upa1 (Fig. 7B). Both proteins, Vps8 and Upa1 have endosomal counterparts in *S. cerevisiae*, namely RING finger E3 ligases Vps8p and Pip1p, respectively (Vps8p and Pib1p, 15, 47).

As pointed out above, Vps8 interacts with the endosomal marker GTPase Rab5 (38), indicating a common theme in endosomal mRNA attachment. In plants, two RRM-type RNA-binding proteins interact with endosomal component *N*-ethylmaleimide-sensitive factor (NSF) as well as Rab5a (48). However, detailed structural information is currently lacking. In humans, a recent high-resolution structural analysis of the FERRY complex (<u>F</u>ive-subunit <u>E</u>ndosomal <u>R</u>ab5 and <u>R</u>NA/ribosome intermediar<u>Y</u>) revealed that the pentameric complex functions as Rab5 effector (49, 50). The integral subunit Fy2 serves as central binding hub, connecting FERRY complex members and mRNAs to Rab5. The complex exhibits a novel clamp-like structure for RNA binding, involving no classic RNA-binding domain but rather coiled coil regions (49). However, whether cargo mRNAs are recognized in a sequence-specific manner has not been clarified yet. Potential cargo mRNAs encode mitochondrial proteins (50), which is a shared feature with the previously reported mRNA cargos of fungal endosomal transport (11).

Studying the MLLE/PAM2L interaction has provided a detailed understanding of the key components involved in endosomal mRNPs attachment. Our findings align with recent perspectives suggesting that RBPs form intricate interaction networks using SLiMs (20, 21). The PAM2 and related SLiMs are known to play roles in network formation via the MLLE^PABPC1^. Here, we add another layer of complexity by demonstrating that two related MLLE domains form a sophisticated network. On one hand, they share interaction partners and binding sequences, but on the other hand, they recognize specific partners using distinct SLiMs (Figure 7B).

## Conclusion

The hitchhiking of mRNAs with shuttling endosomes represents a common mode of trafficking observed across eukaryotes, including fungi, plants and humans. This phenomenon is implicated in a diverse array of processes such as the growth of infectious hyphae, endosperm development and neuronal functions (6, 51, 52). Currently, the most comprehensive understanding exists within fungi, where extensive knowledge on the set of molecular motors (53), the molecular identity of Rab5a-positive endosomes, adaptors, scaffold proteins, key RNA-binding protein, as well as cargo mRNAs with associated ribosomes are available (6). Now, we provide mechanistic insights into the intricate network of RNA-binding proteins during endosomal transport. RBPs rely on MLLE domains recognizing distinct SLiMs in interaction partners. By deciphering the underlying binding code, we have identified new interaction partners in both fungi and humans. Studying fundamental principles of mRNA transport in the microbial model provides a better understanding of pathogenic development (1, 54) and might guide future research endeavors in plant and neuronal systems.

## Materials and methods

### Structure prediction, modelling and analysis

To obtain three dimensional (3D) structural models of the domains and full-length proteins, we utilized Alphafold2 algorithm (29). Monomeric 3D models were generated by providing the protein sequences as input with default parameters in the AlphaFold2_advanced colab notebook (30). Five models were generated for each protein sequence, and the best model was selected based on the pLDDT ranking. Structural analysis and comparison were conducted using the PyMOL molecular graphics system (version 2.0, Schrödinger) and UCSF ChimeraX (version 1.7.1, 55). Interface residues were identified using the PDBePisa server and LigPlot^+^ (56, 57).

### Plasmids, strains, and growth conditions

*E. coli* (K12) Top10 cells (Thermofisher C404010) were utilized for molecular cloning and plasmid DNA propagation, while *E. coli* BL21(DE3) LOBSTR cells (Kerafast EC1002) were employed for recombinant protein expression and purification. Sequences encoding MLLE3^Rrm4^, MLLE^Pab1^ were inserted into the pET22 vector (Merck 69744) with a hexa-histidine tag at the N-terminus (Fig. 1A, H-Rrm4-M3 and H-Pa1-M) for affinity purification. Additionally, sequences encoding MLLE3 variants (wildtype, and mutations, SI Tables S11– S12) were inserted into the pGEX-2T vector (Merck GE28-9546-53) with Glutathione S-Transferase (GST) sequence at N-terminus for pull-down experiments. Sequences encoding PAM2 and PAM2L variants were inserted into the Champion pET-Sumo vector (Thermofisher K30001) with a hexa-histidine and SUMO fusion tag at the N-terminus (HS-PAM2/PAM2L) for the pull-down experiments. Standard techniques were applied for *E. coli* transformation, cultivation, and plasmid isolation.

All *Ustilago maydis* strains are derivatives of AB33 strain, in which hyphal growth is induced by switching the nitrogen source in the medium (33). *U. maydis* yeast-like cells were cultivated in complete medium (CM) supplemented with 1% glucose, while hyphal growth was induced by transferring to nitrate minimal medium (NM) supplemented with 1% glucose. Incubation was carried out at 28°C with constant agitation at 200 rpm (33). Further details regarding growth conditions and general cloning strategies for *U. maydis* can be found elsewhere (58–60). Plasmids were verified by sanger sequencing, and *U. maydis* strains were generated by transforming progenitor strains with linearized plasmids using SspI, or SwaI restriction enzymes. Successful integration of constructs at the desired locus was confirmed by diagnostic PCR, counter-selection between resistance markers, and Southern blot analysis (59). For ectopic integration, plasmids were linearized with SspI, targeted to the ipS locus (61) and selected with carboxin (Cbx). A detailed description of all plasmids, strains, and oligonucleotides is provided in SI Tables S8–S12.

### Recombinant protein expression

Freshly transformed *E. coli* cells were inoculated in 20 ml of expression media. To achieve high-density expression cultures with tight regulation of induction and expression in shake flasks, we formulated a complex media inspired by the principle of Studier’s autoinduction media (14, 62). Glucose was added to the media to prevent the unintended induction and leaky expression of target protein. Phosphate buffer was included to counteract the acidity resulting from glucose metabolism. Additionally, the medium was supplemented with glycerol, nitrogen, sulphur, and magnesium to promote high-density growth. Unlike Studier’s autoinduction media, our formulation does not contain lactose; allowing for induction with IPTG at desired titer and temperature (62; 1.6 % Tryptone, 1% Yeast extract, 50 mM Na_2_HPO_4_, 50 mM KH_2_PO_4_, 25 mM [NH_4_]_2_SO_4_, 0.5% Glycerol, 0.5% Glucose, 2 mM MgSO_4_). Chloramphenicol (34 mg/ml) was consistently added to the expression media to select for plasmid encoding the rare codons. For selection of the expression construct, ampicillin (100 mg/ml), or kanamycin (200 mg/ml) were used. It’s worth noting that the high concentration of kanamycin was employed to prevent unintended resistance caused by high phosphate concentration (62). This optimized media formulation facilitated the growth of high-density cultures, reaching up to OD_600_ = 16 in overnight cultures. We refer to this media as TurboX media for protein expression. The supernatant from the overnight culture was removed by centrifugation at 4°C, 5000 × *g* for 2 minutes. Cell pellets were resuspended in fresh media with an initial OD_600_ of 0.1 and incubated at 37°C with 200 rpm agitation for approximately 2 hours and 30 minutes until the OD_600_ = 1.2. Protein expression was induced by addition of 1 mM IPTG and incubating at 28°C, 200 rpm, for 4 hours, followed by harvesting via centrifugation at 4°C, 6,000 × *g* for 5 minutes.

### Protein purification

For crystallography and ITC experiments, MLLE variants were purified following the methodology outlined in our previous report (14). In brief, the hexa-histidine tagged H-Rrm4-M protein was purified using Nickel-based affinity chromatography (HisTrap HP, GE Healthcare) on Akta primeplus FPLC system (GE Healthcare). Cell pellets were thawed on ice and resuspended in buffer A (20 mM HEPES pH 8.0, 200 mM NaCl, 1 mM EDTA, 10 mM Imidazole pH 8.0, 1 mM PMSF, 0.5 mg/ml Lysozyme, 0.5 mg/ml DNase). Subsequently, cells were lysed by sonication on ice and centrifuged at 4°C, 18,000 × *g* for 30 minutes. The resulting supernatant was loaded onto a pre-equilibrated column with buffer B (20 mM HEPES pH 8.0, 200 mM NaCl,10 mM Imidazole), washed with buffer C (20 mM HEPES pH 8.0, 200 mM NaCl, 50 mM Imidazole), eluted with buffer D (20 mM HEPES pH 8.0, 200 mM NaCl, 300 mM Imidazole), and further purified by size exclusion chromatography (HiLoad 26/600 Superdex 200, GE Healthcare), pre-equilibrated with storage buffer E (20 mM HEPES pH 8.0, 200 mM NaCl). The H-Pab1-M version was purified using the same protocol, with the exception that the wash buffer C was prepared with 20 mM Imidazole (20 mM HEPES pH 8.0, 200 mM NaCl, 20 mM Imidazole). The purity of proteins was assessed by SDS-PAGE. Purified protein samples were concentrated using Amicon 10,000 MWCO centrifugal filter units (Merck, Germany) and stored on ice at 4°C until use. Before use, protein samples were ultra-centrifuged at 4°C, 100,000 × *g* for 30 minutes and quantified by Nanodrop (A280). Peptides were custom-synthesized and purchased from Genscript, USA (see Fig. 1A for peptide sequence).

### GST pull-down experiments

Pull-down assays were conducted following established procedures (14). Briefly, GST-MLLE variants and HS-PAM2/PAM2L variants were expressed in *E. coli* LOBSTR strain (Kerafast EC1002). Cell pellets from 50 ml *E. coli* expression culture were resuspended in 10 ml buffer F (20 mM HEPES pH 8.0, 200 mM NaCl, 1 mM EDTA; 0.5% Nonidet P-40, 1 mM PMSF, 0.1 mg/ml Lysozyme). After sonication on ice, the lysate was centrifuged at 4°C, 16,000 × *g* for 30 minutes. One milliliter of the supernatant was then incubated with 100 µL Glutathione Sepharose (GS) resin (GE Healthcare), pre-equilibrated in buffer F for 1 hour at 4°C with constant agitation at 1,000 rpm. The GS resin was washed three times with 1 ml of buffer G (20 mM HEPES pH 8.0, 200 mM NaCl, 1 mM EDTA, 0.5 % Nonidet P-40). Subsequently, the supernatant containing HS-PAM2/PAM2L variants was added to the GST-MLLE variant bound resins and incubated for 1 hour at 4°C with agitation. Following the incubation, the resins were washed as aforementioned, resuspended in 50 µL of 4x Laemmli loading buffer and 50 µL of buffer G, and boiled for 15 minutes at 95 °C. 25 µL of the sample were loaded onto 12% SDS PAGE gels for analysis, followed by western blotting was using anti-His primary antibody (Sigma-Aldrich, H1029) and anti-mouse IgG HRP conjugate (Promega, W4021) as the secondary antibody. Detection was performed using ECLTM Prime (Cytiva, GERPN2236). Images were captured using the ImageQuant^TM^ LAS4000 luminescence image analyzer, (GE Healthcare) in accordance with the manufacturer’s instructions.

### Crystallization of MLLE3^Rrm4^ and MLLE^Pab1^

IMAC-purified MLLE versions were utilized for co-crystallization studies. Synthetic PAM2^Upa1^ and PAM2L1,2^Upa1^ peptides were dissolved in storage buffer E (20 mM Hepes pH 8.0, 200 mM NaCl). Prior to use, protein samples were centrifuged at 100,000 × *g* for 30 minutes and quantified by Nanodrop (A280), then mixed with the PAM2^Upa1^ or PAM2L1,2^Upa1^ peptide variant in a 1:1.5 molar ratio to achieve a final concentration of 12 mg/ml. Initial crystallization conditions were screened using MRC-3, 96-well sitting drop plates, and various commercially available crystallization screening kits at 12 °C. A volume of 0.1 µL homogeneous protein-peptide solution was mixed with 0.1 µL reservoir solution and equilibrated against 40 µL of the reservoir. After one-week, initial rod-shaped crystals were found which were further optimized by slightly varying the precipitant concentrations. Optimization was also conducted in sitting drop plates (24-well) at 12°C but by mixing 1 µL protein solution with 1 µL of the reservoir solution, equilibrated against 300 µL reservoir solution. The best diffracting crystals of MLLE3^Rrm4^ with PAM2L1^Upa1^ and MLLE3^Rrm4^ with PAM2L2^Upa1^ complexes were grown within 7 days in 0.1 M Sodium HEPES pH 7.5, 25% PEG 3000. The best diffracting crystals of MLLE^Pab1^with PAM2 complex were grown within 7 days in 3.2 M AmSO_4_, 0.1M MES pH 6. Before harvesting the crystals, crystal-containing drops were overlaid with 2 µL mineral oil and immediately frozen in liquid nitrogen.

### Data collection, processing, and structure refinement

The complete diffraction data set of the MLLE complexes (H-Rrm4-M3 with PAM2L1^Upa1^, H-Rrm4-M3 with PAM2L2 ^Upa1^, H-Pab1-M with PAM2^Upa1^) were collected at beamline ID23EH1 in Hamburg, Germany at 100 K and wavelength 0.98 Å, achieving resolutions up to 2.6 Å. All data underwent processing using the automated pipeline at the EMBL HAMBURG and were subsequently reprocessed using XDS (63). AlphaFold2 predicted models for MLLE3^Rrm4^ and MLLE^Pab1^ successfully phased the 1.7 Å data set of MLLE3^Rrm4^-PAM2L1^Upa1^, 2.4 Å data set of MLLE3^Rrm4^-PAM2L2^Upa1^, 2.0 Å data set of MLLE^Pab1^-PAM2^Upa1^, using the program Phaser from the program Phenix suite (64). The structure was then refined in iterative cycles of manual building and refinement using the program Coot (65), followed by software-based refinements using the Phenix suite (64). All residues were found within the preferred and additionally allowed regions of the Ramachandran plot, detailed data collection and refinement statistics are listed in the SI Table S1.

### Isothermal titration calorimetry

All ITC experiments were conducted following the previous report (14). Prior to use, all the protein samples used underwent centrifugation at 451,000 × *g* for 30 minutes and were quantified by Nanodrop (A280). The concentration of MLLE versions (Fig. 1A, H-Rrm4-M3 and H-Pa1-M) were adjusted to 100 µM while PAM2 peptide variants were adjusted to 1200 µM using buffer G (20 mM HEPES pH 8.0, 200 mM NaCl). Using an MicroCal iTC200 titration calorimeter (Malvern Panalytical technologies), a peptide variant with a volume of 40 µL was titrated to the different MLLE versions. Each experiment was conducted at least twice, maintaining consistency. ITC measurements were carried out at 25 °C with a total of 40 injections (1 μL each). The initial injection, with a volume of 0.5 μL, was disregarded from the isotherm. Technical parameters included a reference power of 5 μcal s^-1^; a stirring speed of 750 rpm, a spacing time of 120 s, and a filter period of 5 s. The resulting isotherm was analyzed by fitting it with a one-site binding model using MicroCal ITC-Origin (Microcal LLC).

### Microscopy, image processing and image analysis

Laser-based epifluorescence microscopy was conducted using a Zeiss Axio Observer.Z1, following previous report (14).To assess uni– and bipolar hyphal growth, cells were cultured in 30 ml volumes until reaching an OD_600_ of 0.5, after which hyphal growth was induced. After 6 hours, more than 150 hyphae per strain were examined for growth behavior (n = 3). Cells were scored for unipolar and bipolar growth, and for the formation of a basal septum. For the analysis of signal number, velocity and distance traveled by fluorescence-labeled Rrm4-Kat variants, movies were recorded with an exposure time of 150 ms and 150 frames. Over 20 hyphae were analyzed per strain (n = 3). All movies and images were processed and analyzed using Metamorph software (version 7.7.0.0, Molecular Devices, Seattle, IL, USA). For both micrographs and kymographs, a segment of 20 µm from the hyphal tip was utilized. To statistically analyze the signal number, velocity and distance travelled, processive signals covering a distance of more than 5 µm were manually counted. All collected data points are depicted, with individual replicates represented in various shades of grey for clarity, while mean values are highlighted in red. Two-tailed Student’s t-tests were employed for all statistical analyses.

### Data, Materials and Software Availability

The X-ray crystallographic data have been deposited in Protein Data Bank (https://www.rcsb.org) under accession number (PDB ID: 8S6N, 8S6O and 8S6U) for complexes MLLE3^Rrm4^ with PAM2L1^Upa1^, MLLE3^Rrm4^ with PAM2L2^Upa1^ and MLLE^Pab1^ with PAM2^Upa1^, respectively. Strains, plasmids and their sequences are available upon request. All other data are included in the manuscript and/or supporting information.

## Acknowledgements

We thank laboratory members for critically reading the manuscript. We acknowledge Drs. Georg Groth and Lutz Schmitt for supporting us with ITC experiments. We thank Dr. Astrid Port, Violetta Applegate and Stefanie Galle from Center for Structural studies (CSS) at HHU for support with X-ray crystallization. The synchrotron MX data were collected at ESRF ID23-EH1. We thank Sylvain Engilberge for the assistance in using the beamline. We are grateful to Dihia Moussaoui at the ESRF for providing assistance during data collection. We acknowledge Drs. Cornelia Rücklé, Kathi Zarnack and Julian König for the insights on the PAM2 sequences of Makorin. The work was funded by grants from the Deutsche Forschungsgemeinschaft under Germany’s Excellence Strategy EXC-2048/1 – Project ID 39068111 to MF; Project-ID 267205415 – SFB 1208 to MF (project A09); Project ID 458090666 – SFB1535 to MF (project A03), and FA (project B02) and SHJS (project Z01). The CSS is funded by the Deutsche Forschungsgemeinschaft (DFG Grant number 417919780; INST 208/740-1 FUGG; INST 208/868-1 FUGG).

## Author contributions

SKD, SS, FA and MF designed this study and analyzed the data. SKD and SS contributed equally to the structural biology and biochemistry. KM and SS performed the cell biology experiments. KM and SKD coordinated strain generation and experimental design. SHJS, SKD and FA contributed to X-ray structure analysis. SKD and MF drafted and revised the manuscript with input from all co-authors. MF contributed funding and resources.

## Expanded View Figures

**Figure EV1.**
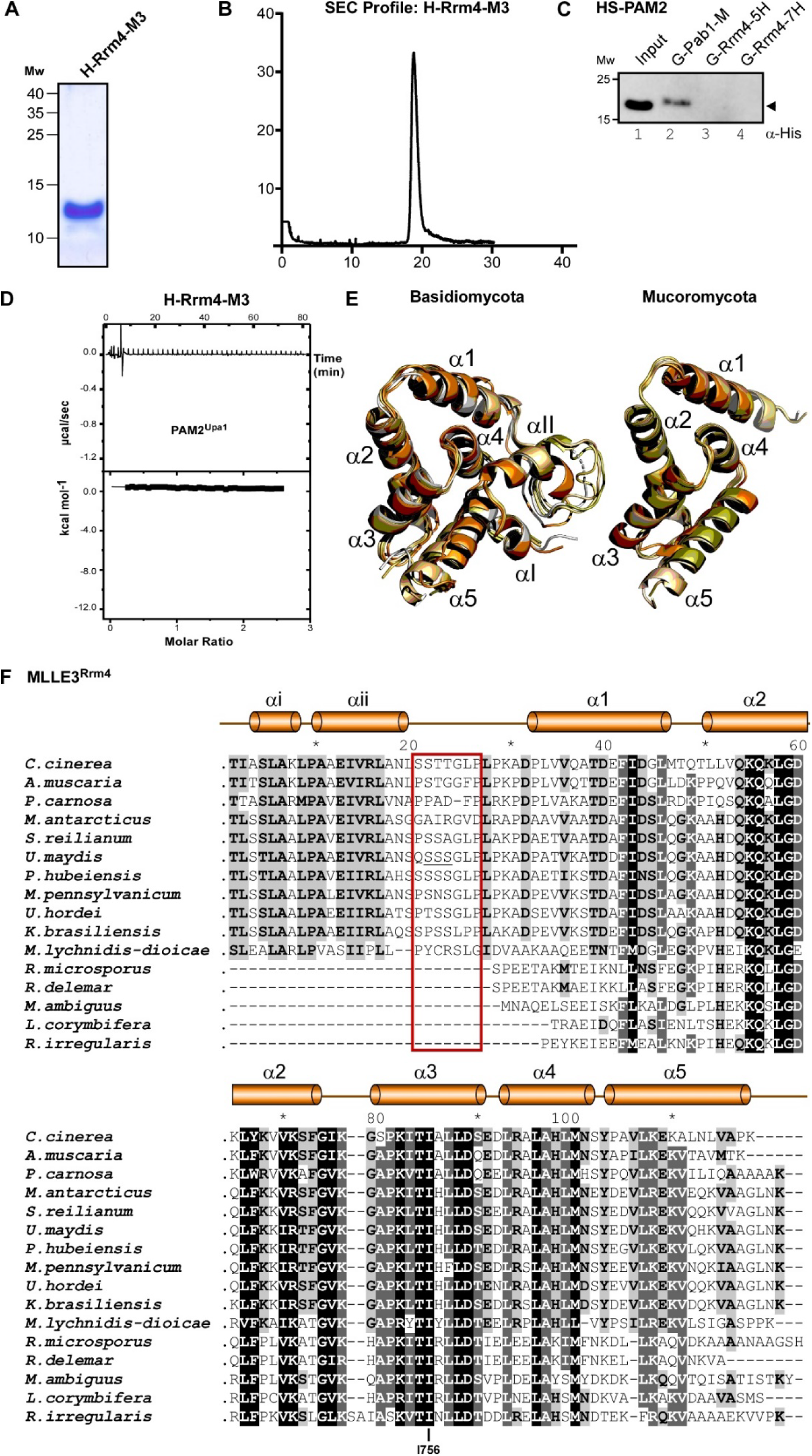
The seven-helix type MLLE domain is evolutionarily conserved. (**A**) SDS-PAGE analysis of purified H-Rrm4-M3 used in crystallography and ITC. (**B**) SEC analysis of purified H-Rrm4-M3. (**C**) Western blot analyses of GST pull-down experiments using α-His for detection, (input, respective His_6_-SUMO peptides). (**D**) ITC binding curves of MLLE3^Rrm4^ domain (H-Rrm4-M3) with PAM2^Upa1^. No interaction detected. (**E**) Overlay of Alphafold predicted models of MLLE3^Rrm4^ orthologues from Basidiomycota consisted of 7 helices (left; grey, *U. maydis;* yellow orange, S. *reilianum;* deep olive, *C. cinerea;* light orange, *P. carnosa;* Orange, *M. lychnidis-dioicae*), Mucoromycota consisted of 5 helices (right; wheat, *R. microspores*, yellow orange, *R. delemar*, light orange, *M. ambiguus*, deep olive, *L. corymbifera*, Orange, *R. irregularis*). (**F**) Multiple sequence alignment of MLLE3^Rrm4^ orthologs of representative fungi from Basidiomycota (*C. cinerea; A. muscaria, P. carnosa, M. antarcticus, S. reilianum, U. maydis, P. hubeiensis, M. pennsylvanicum, U. hordei, K. brasiliensis, M. lychnidis-dioicae*) and Mucoromycota (pale yellow, *R. microspores;* wheat*, M. ambiguus,* light orange, *R. delemar*, orange*, L. corymbifera;* olive, *R. irregularis*). Accession number and sequence coverage are listed in SI Table S2. Red box indicates the serine/threonine rich linker.

**Figure EV2.**
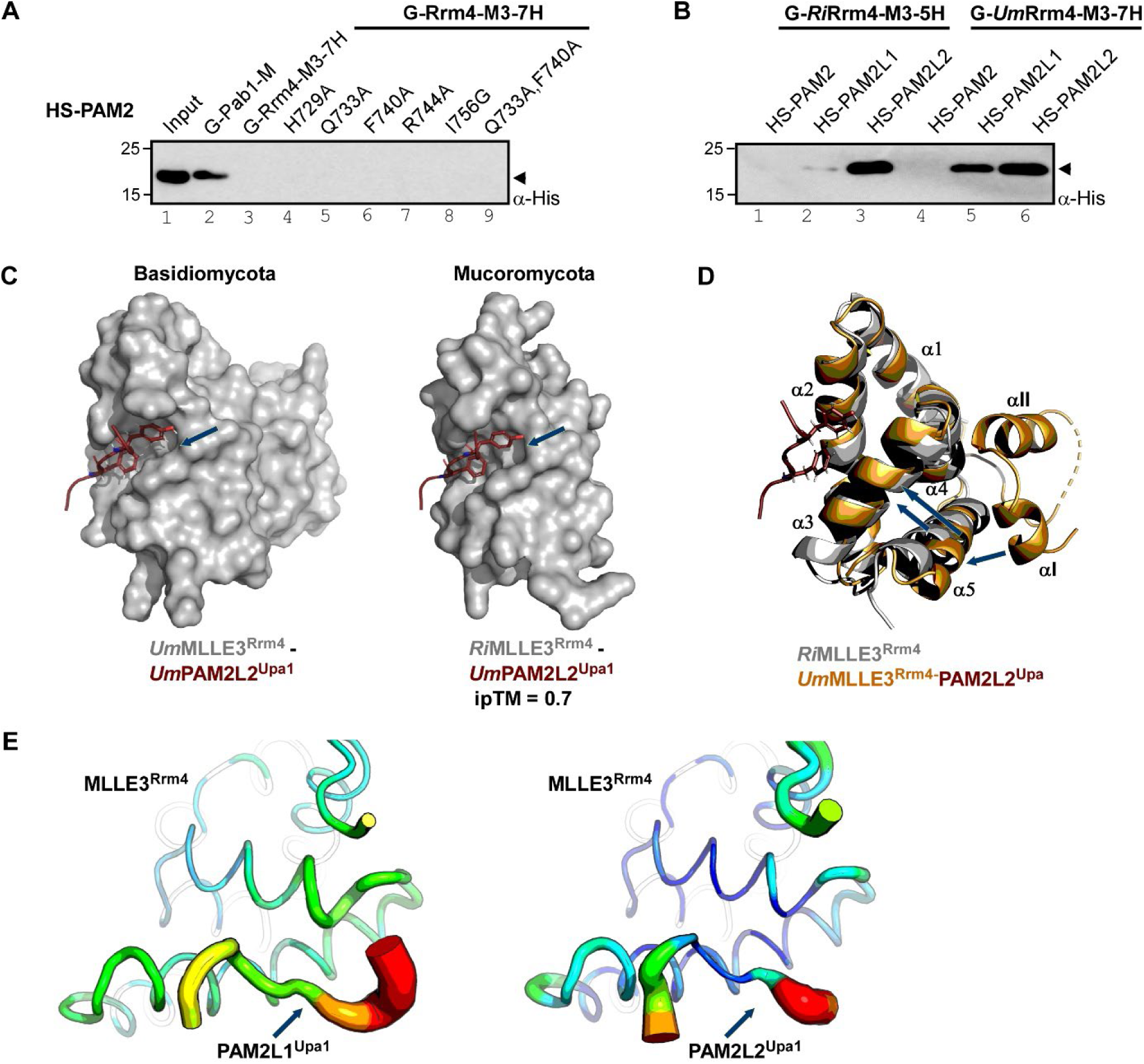
Comparison of seven-helix and five-helix versions of MLLE3^Rrm4^ domains. (**A-B**) Western blot analysis of GST pull-down experiments using α-His for detection (input, respective His_6_-SUMO peptides). (**C**) Comparison of co-crystallized complex of MLLE3^Rrm4^ from *U. maydis* (left, grey surface) consisted of seven-helices bound to PAM2L2^Upa1^ (red sticks) with Alphafold predicted complex of MLLE3^Rrm4^ from *R. irregularis* (right, grey surface) consisted of five-helices bound to PAM2L2^Upa1^ (red sticks) (**D**) Overlay of five and seven-helix-type versions of MLLE3 from *U. maydis* (orange cartoon) and *R. irregularis* (grey cartoon) (**E**) Local displacement of atoms within the MLLE3^Rrm4^-PAM2L1^Upa1^ (left) and MLLE3^Rrm4^-PAM2L1^Upa1^ (right) complexes indicating flexible versus rigid regions as suggested by b-factors (red to blue: high b-factors to low b-factors).

**Figure EV3.**
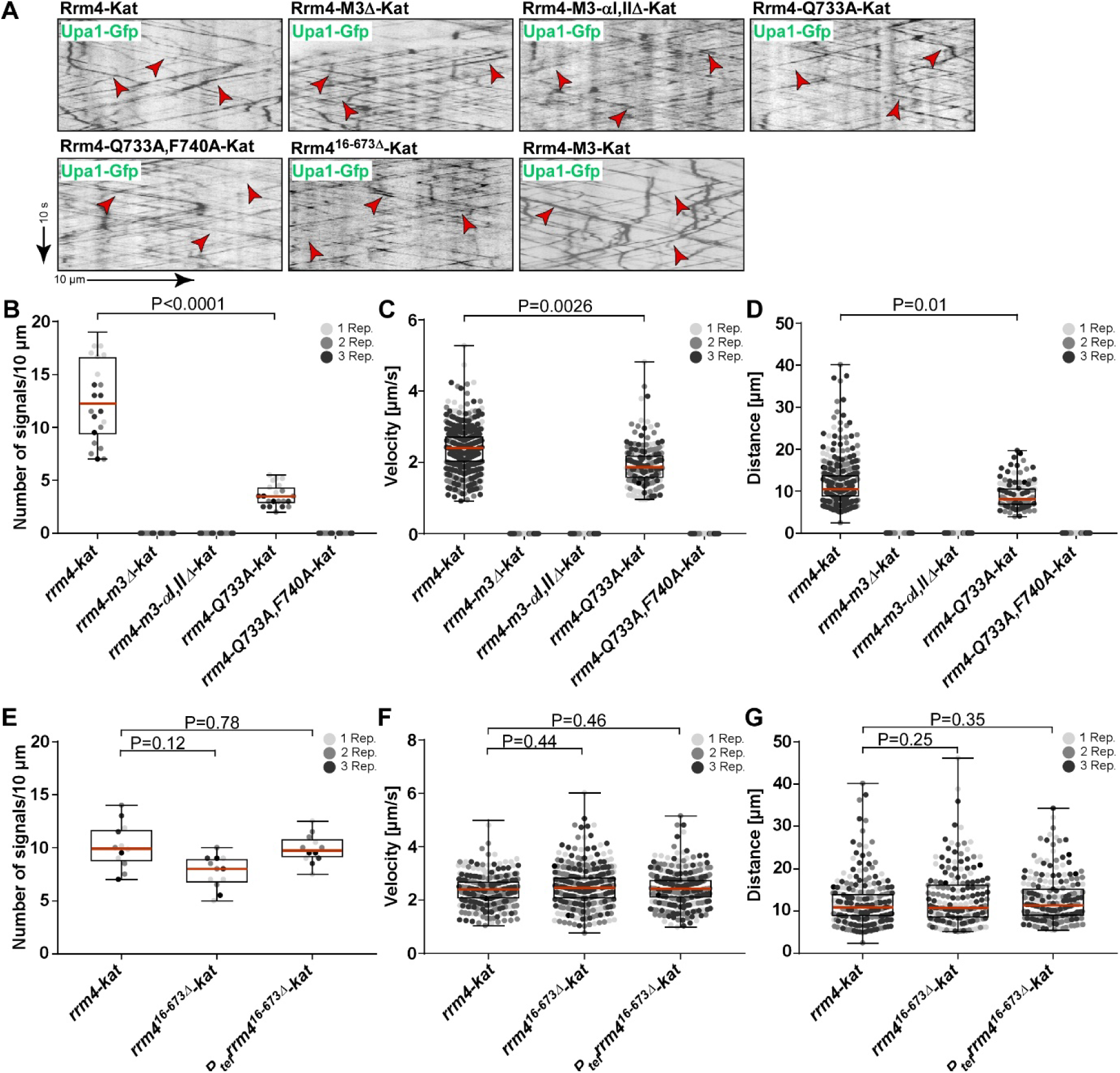
Loss of endosomal localization of Rrm4 caused by the absence of αi and αii helices and the mutation of key amino acids QF740 and Q733 in MLLE3^Rrm4^. (**A**) Kymographs of AB33 hyphae derivatives (6 h.p.i.) showing movement of Upa1-Gfp in hyphae coexpresing different Rrm4-Kat versions (inverted fluorescence images; arrow length on the left and bottom indicates time and distance, respectively). Processive signals are indicated by red arrowheads. (**B-G**) Quantification of processive Rrm4-Kat signals (B and E), velocity of fluorescent Rrm4-Kat signals (C and F) and the traveled distance of processive Rrm4-Kat (D and G; exemplarily kymographs are shown in figure 3; per 10 μm of hyphal length; only particles with a processive movement of > 5 μm were conducted; the three replicates are shown in different gray levels for better identification, red line shows median, SEM; unpaired two-tailed Student’s t-test (α<0.05), for each experiment more than 20 hyphae were analyzed per strain).

**Figure EV4.**
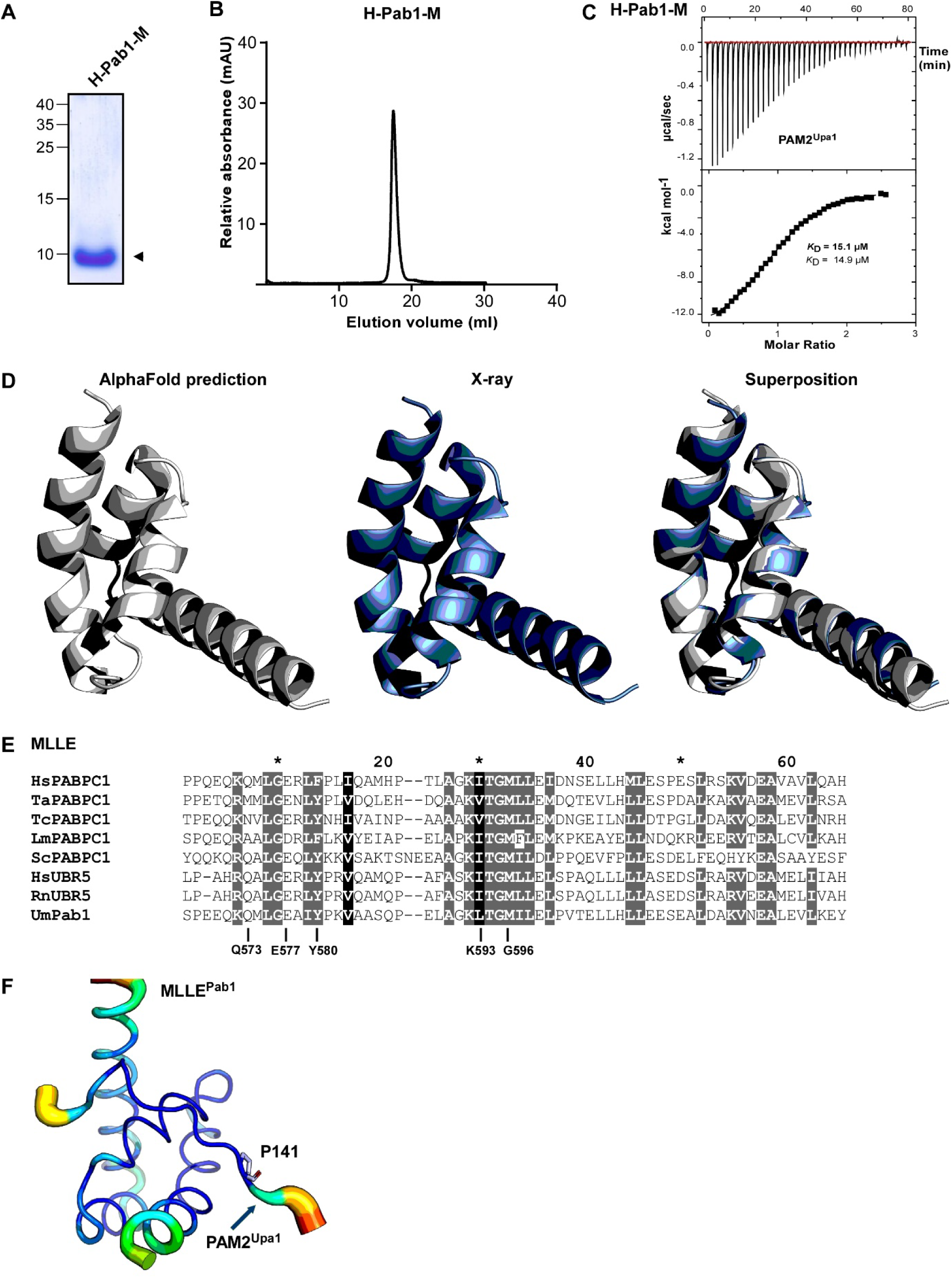
The MLLE domain of Pab1 exhibits a classical α-helical bundle. (**A**) SDS-PAGE analysis of purified H-Pab1-M used in crystallography and ITC. (**B**) SEC analysis of purified H-Pab1-M. (**C**) Representative ITC binding curve of MLLE^Pab1^ domain (H-Pab1-M) with ligand PAM2^Upa1^. KD values of two independent measurements are given (indicated data in bold). (**D**) Structural models of MLLE^Pab1^, generated using Alphafold (grey), X-ray (blue) and overlay as indicated. (**E**) Comparison of MLLE^Pab1^ sequence with structure determined orthologs (Hs – *Homo sapiens*, Ta – *Triticum aestivum*, Tc –*Trypanosoma cruzi*, Lm – *Leishmania major*, Sc – *Saccharomyces cerevisiae*, Rn – *Rattus norvegicus*, Um – *Ustilago maydis*, PABPC1, Pab1 –poly [A]-binding protein, UBR5 – E3 ubiquitin-protein ligase). Accession number and sequence coverage are listed in SI Table S3 (**F**) Local displacement of atoms within the MLLE^Pab1^-PAM2^Upa1^ complex indicating flexible versus rigid regions as suggested by b-factors are shown (red to blue: high b-factors to low b-factors).

**Figure EV5.**
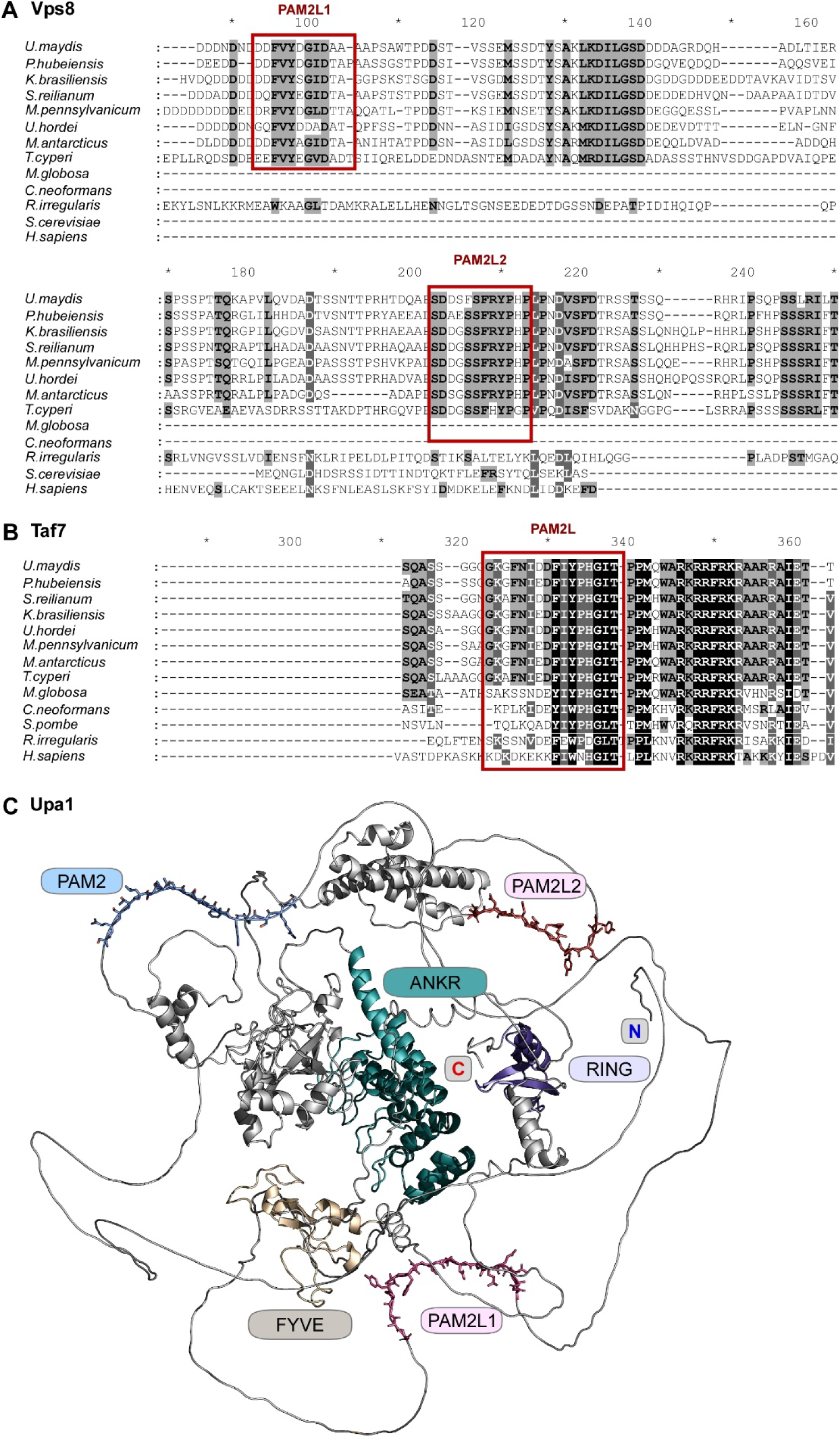
De novo predicted interaction partners of MLLE3^Rrm4^ and Alphafold predicted model of Upa1. Multiple sequence alignment of Vps8 (**A**) and Taf7 (**B**) orthologs *(Ustilago maydis*, *Pseudozyma hubeiensis, Kalmanozyma brasiliensis, Sporisorium reilianum, Ustilago hordei, Moesziomyces pennsylvanicum, M. antarcticus, Testicularia cyperi, Malassezia globosa, Cryptococcus neoformans var. grubii, Rhizophagus irregularis, Saccharomyces cerevisiae, Schizosaccharomyces pombe, Homo sapiens,* accession numbers are listed in the SI Table S5-S6). PAM2L sequences are denoted by red box. (C) Structural model of Upa1 predicted using Alphafold indicating the presence of PAM2^Upa1^ (blue sticks) and PAM2L1,2^Upa1^ (Pink sticks) motifs at the intrinsically disordered region.

**Figure EV6.**
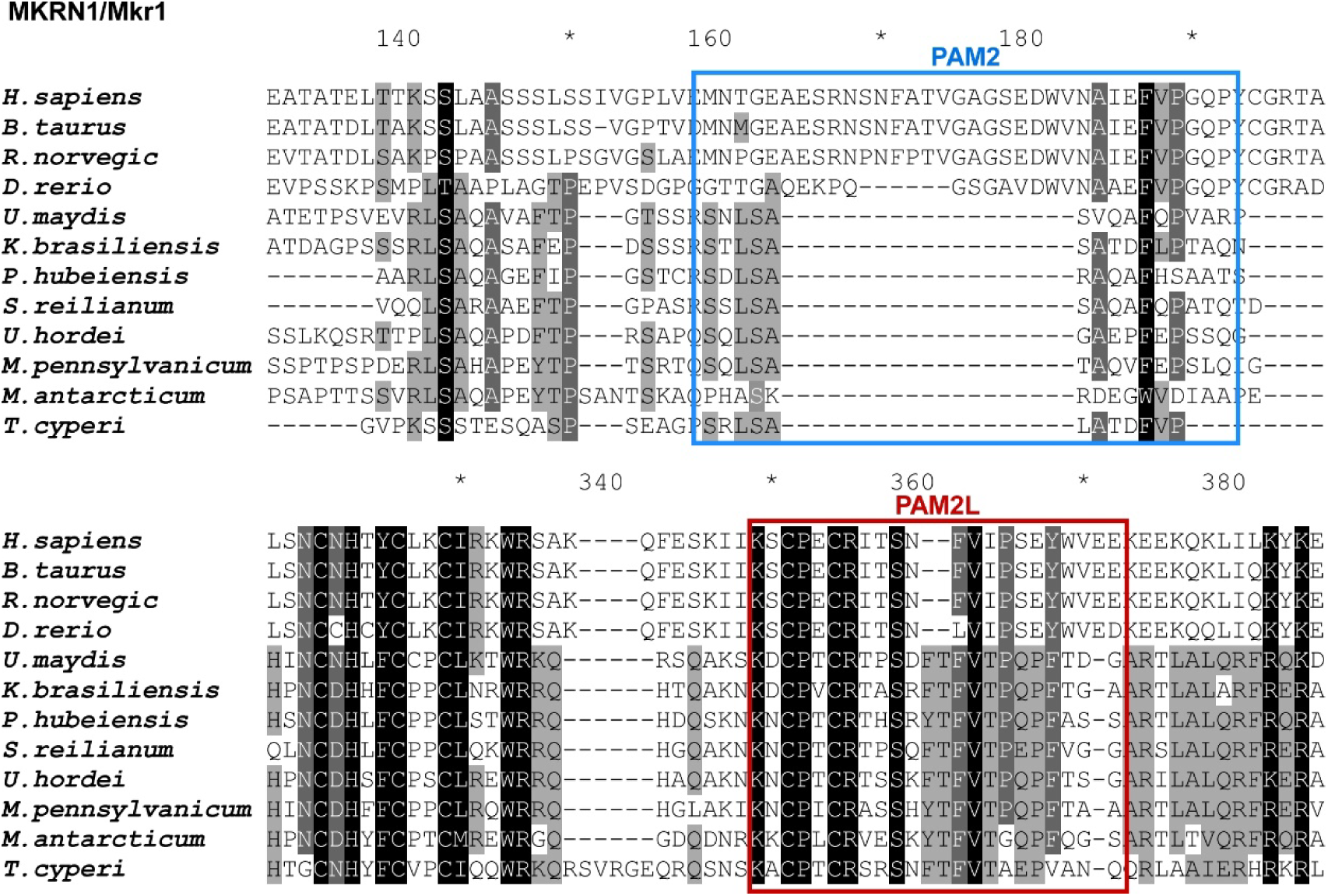
Makorin from human and *U. maydis* contain PAM2L sequences. Multiple sequence alignment of MRKN1 (Human) and Mkr1 (*U. maydis*) orthologs. PAM2, PAM2L sequences are indicated by blue and red boxes respectively.

