## Supplemental Information for "Deciphering the RNA-binding protein network during endosomal mRNA transport"

$ shared first authorship

[1] Institute of Microbiology, Heinrich Heine University Düsseldorf, Cluster of Excellence on Plant Sciences, 40204 Düsseldorf, Germany
[2] Center for Structural Studies, Heinrich Heine University Düsseldorf, 40204 Düsseldorf, Germany
[3] Institute of Biochemistry, Heinrich Heine University Düsseldorf, 40204 Düsseldorf, Germany

#### Table S1. Data collection and refinement statistics for X-ray structures

| Parameters | MLLE3$^{Rrm4}$-PAM2L1$^{Upa1}$ | MLLE3$^{Rrm4}$-PAM2L2$^{Upa1}$ | MLLE$^{Pab1}$-PAM2$^{Upa1}$ |
|---|---|---|---|
| Wavelength | 0.9253 | 1 | 0.8856 |
| Resolution range | 46.61 - 1.743 (1.805 - 1.743) | 56.17 - 2.4 (2.486 - 2.4) | 40.26 - 2.0 (2.072 - 2.0) |
| Space group | C 2 2 21 | C 1 2 1 | C 1 2 1 |
| Unit cell | 74.715 83.486 170.398 90 90 90 | 82.881 74.633 168.511 90 90.081 90 | 88.093 47.608 45.452 90 117.667 90 |
| Total reflections | 224109 (17004) | 120083 (11963) | 22019 (1893) |
| Unique reflections | 54171 (4939) | 39386 (3912) | 11064 (953) |
| Multiplicity | 4.1 (3.4) | 3.0 (3.1) | 2.0 (2.0) |
| Completeness (%) | 98.91 (91.55) | 97.17 (98.27) | 97.13 (85.53) |
| Mean I/sigma (I) | 9.06 (0.86) | 8.20 (1.40) | 18.81 (3.13) |
| Wilson B-factor | 26.97 | 49.66 | 34.86 |
| R-merge | 0.08617 (1.193) | 0.09393 (1.046) | 0.02212 (0.2309) |
| R-meas | 0.09871 (1.409) | 0.1134 (1.261) | 0.03128 (0.3265) |
| R-pim | 0.04717 (0.73) | 0.06273 (0.6948) | 0.02212 (0.2309) |
| CC1/2 | 0.998 (0.361) | 0.997 (0.581) | 0.999 (0.875) |
| CC* | 0.999 (0.728) | 0.999 (0.857) | 1 (0.966) |
| Reflections used in refinement | 54087 (4938) | 39281 (3912) | 11055 (952) |
| Reflections used for R-free | 2667 (232) | 897 (93) | 1100 (95) |
| R-work | 0.2200 (0.3471) | 0.2399 (0.3498) | 0.2017 (0.2289) |
| R-free | 0.2498 (0.3993) | 0.2760 (0.4326) | 0.2524 (0.3182) |
| CC (work) | 0.944 (0.628) | 0.923 (0.644) | 0.950 (0.878) |
| CC (free) | 0.921 (0.592) | 0.900 (0.261) | 0.962 (0.876) |
| Number of non-hydrogen atoms | 4087 | 7457 | 1363 |
| Macromolecules | 3816 | 7424 | 1254 |
| Solvent | | | 10 |
| Protein residues | 271 | 33 | 99 |
| RMS(bonds) | 504 | 968 | 163 |
| RMS(angles) | 0.008 | 0.025 | 0.008 |
| Ramachandran favored (%) | 1.30 | 2.34 | 1.01 |
| Ramachandran allowed (%) | 96.52 | 96.69 | 99.35 |
| Ramachandran outliers (%) | 2.87 | 2.46 | 0.65 |
| Rotamer outliers (%) | 0.61 | 0.85 | 0.00 |
| Clashscore | 0.00 | 0.12 | 6.3 |
| Average B-factor | 6.89 | 18.74 | 2.76 |
| Macromolecules | 33.35 | 65.37 | 39.36 |
| Solvent | 32.93 | 65.42 | 42.19 |

Statistics for the highest-resolution shell are shown in parentheses

#### Table S2. Accession numbers for protein sequences used in multiple sequence alignment of MLLE3-type domain from Rrm4 orthologus

| Organism Name | Fungal phylum | Uniprot/NCBI ID | Sequence coverage |
|---|---|---|---|
| *Coprinopsis cinerea* | Basidiomycota | A8NCM2 | 751-862 |
| *Amanita muscaria* | Basidiomycota | A0A0C2SHU5\| | 675-786 |
| *Phanerochaete carnosa* | Basidiomycota | XP_007393387.1 | 672-785 |
| *Moesziomyces antarcticus* | Basidiomycota | XP_014657015.1 | 684-798 |
| *Sporisorium reilianum* | Basidiomycota | CBQ73718.1 | 660-785 |
| *Ustilago maydis* | Basidiomycota | A0A0D1DWZ5 | 752 - 792 |
| *Pseudozyma hubeiensis* | Basidiomycota | XP_012192836.1 | 537-651 |
| *Melanopsichium pennsylvanicum* | Basidiomycota | CDI54139.1 | 687-800 |
| *Ustilago hordei* | Basidiomycota | I2FYN4 | 685-798 |
| *Kalmanozyma brasiliensis* | Basidiomycota | XP_016289934.1 | 573-686 |
| *Microbotryum lychnidis-dioicae* | Basidiomycota | KDE02990.1 | 551-661 |
| *Rhizopus microspores* | Mucoromycota | ORE17079.1 | 553-642 |
| *Rhizopus delemar* | Mucoromycota | I1CQR1 | 487-567 |
| *Mucor ambiguus* | Mucoromycota | GAN10032.1 | 888-974 |
| *Lichtheimia corymbifera* | Mucoromycota | CDH52259.1 | 682-760 |
| *Rhizophagus irregularis* | Mucoromycota | GBC41783.1 | 572-655 |

#### Table S3. Accession numbers for protein sequences used in multiple sequence alignment of MLLE domain in PABPC1/Pab1 and Ubr5 orthologus

| Organism Name | Protein Name | Domain name | Uniprot/NCBI ID |
|---|---|---|---|
| *Homo sapiens* | Poly[A] binding protein, PABP | $MLLE^{PABP}$ | P11940 |
| *Triticum aestivum* | Poly[A] binding protein, PABP | $MLLE^{PABP}$ | P93616 |
| *Trypanosoma cruzi* | Poly[A] binding protein, PABP | $MLLE^{PABP}$ | Q27335 |
| *Leishmania major* | Poly[A] binding protein, PABP | $MLLE^{PABP}$ | E9AFX7 |
| *Saccharomyces cerevisiae* | Poly[A] binding protein, PABP | $MLLE^{PABP}$ | P04147 |
| *Homo sapiens* | E3 ubiquitin-protein ligase UBR5 | $MLLE^{Ubr5}$ | O95071 |
| *Rattus norvegicus* | E3 ubiquitin-protein ligase UBR5 | $MLLE^{Ubr5}$ | Q62671 |
| *Ustilago maydis* | Poly[A] binding protein, Pab1 | $MLLE^{Pab1}$ | Q4P8R9 |

#### Table S4. De novo predicted PAM2L candidates in *U. maydis*

| UMAG | Protein Name | Region | PAM2L sequence |
|---|---|---|---|
| UMAG_15064 | Vps8, early endosome specific subunit of CORVET complex | IDR* | HQDDDNDND**DDF**V**Y**DGID, TDQAPSDDSFS**F**R**Y**PHPL |
| UMAG_02262 | Transcription factor CBF | IDR | LNDSDDDDD**DEF**D**Y**IDSD |
| UMAG_04936 | FCP1 domain-containing protein, that dephosphorylates the C-terminal domain (CTD) of RNA polymerase II | IDR | HSAHGAMAD**DDF**E**Y**DSDL |
| UMAG_04517 | DUF3835 domain protein in fungi | IDR | DQDDFDDQD**DDF**D**Y**DPDD |
| UMAG_01597 | Transcriptional activator HAP2 | IDR | SRAQSVVGS**EDF**A**Y**HASP |
| UMAG_10255 | Orthologue of Yeast transcription corepressor | IDR | HTTPPVAEE**DEF**H**Y**PLPS |
| UMAG_02933 | Uncharacterized protein | IDR | PAAFESLLD**DDF**S**Y**LVDD |
| UMAG_02807 | Methyl-itaconate delta2-delta3-isomerase | IDR | LQPTSSRHF**EDF**R**Y**VEPK |
| UMAG_10091 | Serine carboxypeptidase | IDR | PWRPATVHS**EEF**A**Y**GGNR |
| UMAG_10620 | Taf7, Ptr6 orthologue | Linker | GGGGKGFNI**DDF**I**Y**PHGI |
| UMAG_05811 | GH16 domain-containing protein | Linker | PLLTDQEDD**DDF**L**Y**SDKG |
| UMAG_10591 | Ankyrin repeat domain containing protein | Linker | GEGISPVAA**DEF**T**Y**TPLH |

* IDR – Intrinsically disordered region

#### Table S5. Accession numbers for protein sequences used in multiple sequence alignment of Vps8 orthologs

| Organism Name | Target sequence | Uniprot/NCBI ID |
|---|---|---|
| *Ustilago maydis* | PAM2L1,2 | A0A0D1DXQ1 |
| *Pseudozyma hubeiensis* | PAM2L1,2 | R9P7V6 |
| *Kalmanozyma brasiliensis* | PAM2L1,2 | EST05200.2 |
| *Sporisorium reilianum* | PAM2L1,2 | A0A2N8UEF7 |
| *Melanopsichium pennsylvanicum* | PAM2L1,2 | CDI53747.1 |
| *Ustilago hordei* | PAM2L1,2 | XP_041410257.1 |
| *Moesziomyces antarcticus* | PAM2L1,2 | XP_014656004.1 |
| *Testicularia cyperi* | PAM2L1,2 | PWY99436.1 |
| *Malassezia globosa* | PAM2L1,2 | A8QA38 |
| *Cryptococcus neoformans var grubii* | PAM2L1,2 | J9VWP5 |
| *Rhizophagus irregularis* | PAM2L1,2 | A0A2I1GF06 |
| *Saccharomyces cerevisiae* | PAM2L1,2 | P39702 |
| *Homo sapiens* | PAM2L1,2 | Q8N3P4 |

#### Table S6. Accession numbers for protein sequences used in multiple sequence alignment of Taf7 orthologus

| Organism Name | Target sequence | Uniprot/NCBI ID |
|---|---|---|
| *Ustilago maydis* | PAM2L | XP_011387353.1 |
| *Pseudozyma hubeiensis* | PAM2L | XP_012192603.1 |
| *Sporisorium reilianum* | PAM2L | SJX60978.1 |
| *Kalmanozyma brasiliensis* | PAM2L | EST10146.2 |
| *Ustilago hordei* | PAM2L | XP_041415606.1 |
| *Melanopsichium pennsylvanicum* | PAM2L | CDI51836.1 |
| *Moesziomyces antarcticus* | PAM2L | XP_014658645.1 |
| *Testicularia cyperi* | PAM2L | PWZ00942.1 |
| *Malassezia globosa* | PAM2L | XP_001732308.1 |
| *Cryptococcus neoformans var grubii* | PAM2L | OXB39574.1 |
| *Saccharomyces pombe* | PAM2L | O13701 |
| *Saccharomyces cerevisiae* | PAM2L | Q05021 |
| *Rhizophagus irregularis* | PAM2L | PKC16038.1 |
| *Homo sapiens* | PAM2L | Q15545 |

#### Table S7. Accession numbers for protein sequences used in multiple sequence alignment of MKRN1/Mkr1 orthologus

| Organism Name | Target sequences | Uniprot/*NCBI* ID |
|---|---|---|
| *Homo sapiens* | PAM2, PAM2L | Q9UHC7 |
| *Bos taurus* | PAM2, PAM2L | R9P7K9 |
| *Rattus norvegicus* | PAM2, PAM2L | *NP_001385662.1* |
| *Danio rerio* | PAM2, PAM2L | Q4VBT5 |
| *Ustilago maydis* | PAM2, PAM2L | A0A0D1E4Z6 (UMAG_12122) |
| *Kalmanozyma brasiliensis* | PAM2, PAM2L | *KAF6767369.1* |
| *Pseudozyma hubeiensis* | PAM2, PAM2L | *R9P7K9* |
| *Sporisorium reilianum* | PAM2, PAM2L | *A0A2N8U8L2* |
| *Ustilago hordei* | PAM2, PAM2L | *XP_041415531.1* |
| *Rhizophagus irregularis* | PAM2, PAM2L | *A0A2I1EXN5* |
| *Melanopsichium pennsylvanicum* | PAM2, PAM2L | A0A077R4S2 |
| *Moesziomyces antarcticus* | PAM2, PAM2L | GAC72811.1 |
| *Testicularia cyperi* | PAM2, PAM2L | *PWZ01012.1* |

#### Table S8. Description of *U. maydis* strains used in this study

| Strain name with code | Locus | Progenitor strain | Short description |
|---|---|---|---|
| AB33 (UMa133) | *b* | FB2 | *Pnar:bW2bE1*, expression of active b heterodimer under control of the *nar1* promoter, strain grows filamentous upon changing the nitrogen source. |
| AB33rrm4Δ/upa1-gfp (UMa2769) | *rrm4* *upa1* | AB33rrm4-Cherry/ upa1-gfp | carrying a deletion of *rrm4* and expressing Upa1 C-terminally fused to eGfp |
| AB33upa1-gfp/rrm4-kat (UMa2976) | *rrm4* *upa1* | AB33rrm4Δ/upa1-gfp | expressing Upa1 C-terminally fused to eGfp and Rrm4 C-terminally fused to mKate2 |
| AB33upa1-gfp/rrm4-m3Δ-kat (UMa2979) | *rrm4* *upa1* | AB33rrm4Δ/upa1-gfp | expressing Upa1 C-terminally fused to eGfp and Rrm4-M3Δ C-terminally fused to mKate2. Like rrm4-kat but carrying the deletion of 3rd MLLE domain. Residues of Rrm4 from 689 to 792 were replaced with a HAtag-HRV3C protease recognition site. |
| AB33upa1-gfp/rrm4-m3-αi,iiΔ-kat (UL167) | *rrm4* *upa1* | AB33rrm4Δ/upa1-gfp | expressing Upa1 C-terminally fused to eGfp and Rrm4-M3-αi,iiΔ C-terminally fused to mKate2. Like Rrm4-kat but carrying the deletion of 1$^{st}$and2$^{nd}$ α-helix of MLLE3 domain. Residues of Rrm4 from 678 to 704 were replaced with a HAtag-HRV3C protease recognition site. |
| AB33upa1-gfp/rrm4-Q733A,F740A-kat (UL168) | *rrm4* *upa1* | AB33rrm4Δ/upa1-gfp | expressing Upa1 C-terminally fused to eGfp and Rrm4-Q733A, F740A C-terminally fused to mKate2. Like rrm4-kat but carrying the substitution mutation of Glutamine 733 and Phenylalanine 740 to Alanine of MLLE3 domain. |
| AB33upa1-gfp/rrm4-Q733A-kat (UL169) | *rrm4* *upa1* | AB33rrm4Δ/upa1-gfp | expressing Upa1 C-terminally fused to eGfp and Rrm4-Q733A C-terminally fused to mKate2. Like rrm4-kat but carrying the substitution mutation of Glutamine733 to Alanine of MLLE3 domain. |
| AB33upa1-gfp/rrm4$^{16\text{-}673\Delta}$-kat (UL170) | *rrm4* *upa1* | AB33rrm4Δ/upa1- gfp | expressing Upa1 C-terminally fused to eGfp and Rrm4$^{16\text{-}673\Delta}$ C-terminally fused to mKate2. 658 amino acid residues between position 16 and 673 was deleted from the full-length Rrm4 protein, containing just the MLLE3 domain with 7α-helix |
| AB33upa1-gfp/rrm4/P$_{tef}$ rrm4$^{16\text{-}673\Delta}$-kat (UL171) | *ip$^S$* *upa1* | AB33upa1-gfp | expressing Upa1 C-terminally fused to eGfp, Rrm4$^{16\text{-}673\Delta}$ C-terminally fused to mKate2 ectopically at the *ip$^S$* locus and the Rrm4 at the native locus is undisturbed. 658 amino acid residues between position 16 and 673 was deleted from the full-length Rrm4 protein, containing just the MLLE3 domain with 7α-helix at the *ip$^S$* locus |

#### Table S9. Generation of *U. maydis* strains used in this study

| Strains | Relevant genotype | Strain code | Reference | Transformed plasmid | Locus | Progenitor |
|---|---|---|---|---|---|---|
| AB33 | *a2 $P_{nar}$:bW2 bE1* | UMa 133 | Brachmann, 2001 | pAB33 | *b* | FB2 |
| AB33rrm4Δ/upa1-gfp | *rrm4 upa1-gfp* | UMa 2769 | Devan, 2022 | pRrm4Δ_genitR (pUMa1755) | *rrm4* | AB33rrm4-mCherry/upa1-gfp (UMa1594) |
| AB33upa1-gfp/rrm4-kat | *upa1-gfp rrm4-kat* | UMa 2976 | Devan, 2022 | pRrm4-kat-hygR (pUMa3908) | *rrm4* | AB33rrm4Δ/upa1-gfp (UMa2769) |
| AB33upa1-gfp/rrm4-m3Δ-kat | *upa1-gfp rrm4-m3Δ-kat* | UMa 2979 | Devan, 2022 | pRrm4-m3Δ-kat-hygR (pUMa4435) | *rrm4* | AB33rrm4Δ/upa1-gfp (UMa2769) |
| AB33upa1-gfp/rrm4-m3-αi,iiΔ-kat | *upa1-gfp rrm4-m3-αi,iiΔ-kat* | UL167 | this study | pRrm4- m3- αi,iiΔ-kat-hygR (pUL219) | *rrm4* | AB33rrm4Δ/upa1-gfp (UMa2769) |
| AB33upa1-gfp/rrm4-Q733A,F740A-kat | *upa1-gfp rrm4-Q733A, F740A-kat* | UL168 | this study | pRrm4-Q733A, F740A-kat-hygR (pUL220) | *rrm4* | AB33rrm4Δ/upa1-gfp (UMa2769) |
| AB33upa1-gfp/rrm4-Q733A-kat | *upa1-gfp rrm4-Q733A-kat* | UL169 | this study | pRrm4-Q733A-kat-hygR (pUL222) | *rrm4* | AB33rrm4Δ/upa1-gfp (UMa2769) |
| AB33upa1-gfp/rrm4$^{16\text{-}673\Delta}$-kat | *upa1-gfp rrm4$^{16\text{-}673\Delta}$-kat* | UL170 | this study | pRrm4$^{16\text{-}673\Delta}$-kat-hygR (pUL221) | *rrm4* | AB33rrm4Δ/upa1-gfp (UMa2769) |
| AB33upa1-gfp/rrm4/$P_{tef}$ rrm4$^{16\text{-}673\Delta}$-kat | *upa1- gfp rrm4/ $P_{tef}$ rrm4$^{16\text{-}673\Delta}$-kat* | UL171 | this study | Ptef_pRrm4$^{16\text{-}673\Delta}$-kat-cbxR (pUL254) | *$ip^S$* | AB33upa1-gfp (UMa956) |

#### Table S10. Description of plasmids used for *U. maydis* strain generation

| Plasmid | Plasmid ID | Resistance cassette | Short description |
|---|---|---|---|
| pRrm4Δ | pUMa 1755 | genitR (G418 resistance - SfiI insert of pMF1g) Baumann et al., 2012 | Plasmid vector for generating deletion mutants of *rrm4*. |
| pRrm4-kat-hygR | pUMa 3908 | hygR (Hygromycin resistance - SfiI insert of pMF1h) Brachmann et al., 2004 | Plasmid vector for the expression of Rrm4 C-terminally fused to mKate2. The mKate2 cassette contains the Tnos terminator and the Hyg resistance. The entire coding sequence for the fusion protein is flanked by a 1025 bp upstream region and a 1396 bp downstream region for homologous recombination. |
| pRrm4-m3Δ-kat-hygR | pUMa 4435 | hygR | Plasmid vector for the expression of Rrm4-M3Δ C-terminally fused to mKate2. Like pRrm4-mK-HygR, but carrying the deletion of the 3rd MLLE domain. Residues of Rrm4 from 689-792 were replaced with a HAtag-HRV3C protease recognition site. |
| pRrm4-m3-αi,iiΔ-kat-hygR | pUL 219 | hygR | Plasmid vector for the expression of Rrm4-M3-αi,iiΔ C-terminally fused to mKate2. Like Rrm4-kat but carrying the deletion of 1stand2nd α-helix of MLLE3 domain. Residues of Rrm4 from 678 to 704 were replaced with a HAtag-HRV3C protease recognition site. |
| pRrm4-Q733A, F740A-kat-hygR | pUL 220 | hygR | Plasmid vector for the expression of Rrm4-Q733A, F740A-C-terminally fused to mKate2. Like rrm4-kat but carrying the substitution mutation of Glutamine 733 and Phenylalanine 740 to Alanine of MLLE3 domain. |
| pRrm4-Q733A-kat-hygR | pUL 222 | hygR | Plasmid vector for the expression of Rrm4-Q733A C-terminally fused to mKate2. Like rrm4-kat but carrying the substitution mutation of Glutamine733 to Alanine of MLLE3 domain. |
| pRrm4$^{16\text{-}673\Delta}$-kat-hygR | pUL 221 | hygR | Plasmid vector for the expression of Rrm4$^{16\text{-}673\Delta}$ C- terminally fused to mKate2. 658 aminoacid residues between position 16 and 673 was deleted from the 792 aminoacid long Rrm4 protein, containing just the MLLE3 domain with 7α-helix and initial 15 residues of wildtype Rrm4 |
| Ptef_pRrm4$^{16\text{-}673\Delta}$-kat-cbxR | pUL 254 | cbxR (Carboxin resistance - SfiI insert of pMF2-3c) Brachmann et al., 2004 | Plasmid vector for the ectopic expression of Rrm4$^{16\text{-}673\Delta}$ C-terminally fused to mKate2 under the regulation Tef promoter at the *ip*$^{S}$ locus . 658 aminoacid residues between position 16 and 673 was deleted from the 792 aminoacid long Rrm4 protein, containing just the MLLE3 domain with 7α-helix and initial 15 residues of wildtype Rrm4 |

### Table S11: Description of plasmids used for recombinant expression in *E. coli*

| Plasmid | Plasmid ID | Short description |
|---|---|---|
| pGEX-G-Pab1-M_Um | pUMa 2187 | Plasmid for the expression of the G-Pab1-M. C terminal region of Pab1 comprising amino acid residues from 566 to 651 fused to an N-terminal GST- tag. (Pohlmann et al., eLife 2015) |
| pET28-HS-PAM2$^{Upa1_Um}$ | pUMa 4296 | Plasmid for the expression of the PAM2 motif of Upa1 (SQSTLSPNASVFKPSRS) fused to an N-terminal 6xHis-Sumo tag. (Devan et al., PloS Genetics 2022) |
| pET28-HS_PAM2L1$^{Upa1_Um}$ | pUMa 4297 | Plasmid for the expression of PAM2L1 motif of Upa1 (EAADQEEDQDDFVYPGAD) fused to an N-terminal 6xHis-Sumo tag. (Devan et al., PloS Genetics 2022) |
| pET28-HS-PAM2L2$^{Upa1_Um}$ | pUMa 4298 | Plasmid for the expression of PAM2L2 motif of Upa1 (DEDAADDDDDEFIYPNSY) fused to an N-terminal 6xHis-Sumo tag. (Devan et al., PloS Genetics 2022) |
| pGX-G-Rrm4-M3-5H_Um | pUMa 4701 | Plasmid for the expression of G-Rrm4-M3-5H. MLLE3 domain of Rrm4 comprising amino acid residues from 700 to 792 fused to an N-terminal GST- tag. |
| H-Rrm4-M3 | pUMa 4704 | Plasmid for the expression of Rrm4-M3-7H. MLLE3 domain of Rrm4 comprising amino acid residues from 679 to 792 fused to an N-terminal 6xHis-Sumo tag. |
| H-Pab1-M | pUMa 4794 | Plasmid for the expression of Pab1-M-4H. MLLE domain of Pab1 comprising amino acid residues from 567 to 636AA fused to an N-terminal 6xHis-Sumo tag. |
| pET28-HS_PAM2$^{Tob1_Hs}$ | pUL123 | Plasmid for the expression of PAM2 motif of Human Tob1 (SALSPNAKEFIFPNM) fused to an N-terminal 6xHis-Sumo tag. |
| pET28-HS_PAM2L$^{Ubr5_Hs}$ | pUL124 | Plasmid for the expression of PAM2L motif of Human Ubr5 (DTDDGDDNAPLFYQPGKRGF) fused to an N-terminal 6xHis-Sumo tag. |
| pGX-G-Rrm4-M3-7H_Um | pUL130 | Plasmid for the expression of G-Rrm4-M3-7H. MLLE3 domain of Rrm4 comprising amino acid residues from 679 to 792 fused to an N-terminal GST- tag. |
| pGX-G-Rrm4-M3-7H-Q733A,F740A | pUL131 | Plasmid for the expression of G-Rrm4-M3-7H. MLLE3 domain of Rrm4 comprising amino acid residues from 679 to 792 with two point mutation (Glutamine 733 and Phenylalanine 740 to Alanine) fused to an N-terminal GST- tag. |
| pET28-HS_PAM2L$^{Mkr1_Um}$ | pUL132 | Plasmid for the expression of PAM2L motif of Ustilago Mkr1 (CPTCRTPSDFTFVTPQPF) fused to an N-terminal 6xHis-Sumo tag. Mkr1 (RING-type E3 ubiquitin transferase) |
| pET28-HS_PAM2$^{Mkrn1_Hs}$ | pUL133 | Plasmid for the expression of PAM2 motif of Human MKRN1 (AGSEDWVNAIEFVPGQP) fused to an N-terminal 6xHis-Sumo tag. |
| pET28-HS_PAM2$^{Mkr1_Um}$ | pUL134 | Plasmid for the expression of PAM2 motif of Ustilago Mkr1 (SRSNLSASVQAFQPVAR) fused to an N-terminal 6xHis-Sumo tag. Mkr1 (RING-type E3 ubiquitin transferase) |
| pET28-HS-short-PAM2L1$^{Upa1_Um}$ | pUL138 | Plasmid for the expression of PAM2L1 motif of Upa1 (DDFVYPGAD) fused to an N-terminal 6xHis-Sumo tag. |
| pET28-HS-short-PAM2L2$^{Upa1_Um}$ | pUL139 | Plasmid for the expression of PAM2L2 motif of Upa1 (DEFIYPNSY) fused to an N-terminal 6xHis-Sumo tag. |
| pET28-HS-PAM2v$^{GW182_Hs}$ | pUL146 | Plasmid for the expression of PAM2v motif of Human GW182 (NWPPEFHPGVPWKGLQ) fused to an N-terminal 6xHis-Sumo tag. |
| pET28-HS-PAM2_PAM2L2_hyb | pUL157 | Plasmid for the expression of PAM2 and PAM2L2 hybrid motif of Upa1 (SQSTLSPNADEFIYPNSY) fused to an N-terminal 6xHis-Sumo tag. |

| | | |
|---|---|---|
| pET28-HS_PAM2$^{Paip2_Hs}$ | pUL162 | Plasmid for the expression of PAM2 motif of Human Paip2 (VKSNLNPNAKEFVPGVK) fused to an N-terminal 6xHis-Sumo tag. |
| pET28-HS-PAM2L1-Y250A$^{Upa1}$ | pUL166 | Plasmid for the expression of PAM2L1 motif of Upa1 (EAADQEEDQDDFV**A**PGAD) carrying a point mutation (Tyrosine 250 to Alanine) fused to an N-terminal 6xHis-Sumo tag. |
| pET28-HS-PAM2L2-Y957A$^{Upa1}$ | pUL167 | Plasmid for the expression of PAM2L2 motif of Upa1 (DEDAADDDDDEFI**A**PNSY) carrying a point mutation (Tyrosine 957 to Alanine) fused to an N-terminal 6xHis-Sumo tag. |
| pET28-HS-PAM2L2_PAM2_hyb | pUL168 | Plasmid for the expression of PAM2L2 and PAM2 hybrid motif of Upa1 (DEDAADDDDSVFKPSRS) fused to an N-terminal 6xHis-Sumo tag. |
| pET28-HS-PAM2L1-F248A$^{Upa1}$ | pUL169 | Plasmid for the expression of PAM2L1 motif of Upa1 (EAADQEEDQDD**A**VYPGAD) carrying a point mutation (Phenylalanine 248 to Alanine) fused to an N-terminal 6xHis-Sumo tag. |
| pET28-HS-PAM2L2-F955A$^{Upa1}$ | pUL170 | Plasmid for the expression of PAM2L2 motif of Upa1 (DEDAADDDDDE**A**IYPNSY) carrying a point mutation (Phenylalanine 955 to Alanine) fused to an N-terminal 6xHis-Sumo tag. |
| pET28-HS-PAM2L1-P251A$^{Upa1}$ | pUL171 | Plasmid for the expression of PAM2L1 motif of Upa1 (EAADQEEDQDDFVY**A**GAD) carrying a point mutation (Proline 251 to Alanine) fused to an N-terminal 6xHis-Sumo tag. |
| pET28-HS-PAM2L2-P958A$^{Upa1}$ | pUL172 | Plasmid for the expression of PAM2L2 motif of Upa1 (DEDAADDDDDEFIY**A**NSY) carrying a point mutation (Proline 958 to Alanine) fused to an N-terminal 6xHis-Sumo tag. |
| pET28-HS-PAM2-F139A$^{Upa1}$ | pUL173 | Plasmid for the expression of the PAM2 motif of Upa1 (SQSTLSPNASV**A**KPSRS) carrying a point mutation (Phenylalanine 139 to Alanine) fused to an N-terminal 6xHis-Sumo tag. |
| pET28-HS-PAM2L1-D246A$^{Upa1}$ | pUL174 | Plasmid for the expression of PAM2L1 motif of Upa1 (EAADQEEDQ**A**DFVYPGAD) carrying a point mutation (Aspartate 248 to Alanine) fused to an N-terminal 6xHis-Sumo tag. |
| pET28-HS-PAM2L2-D953A$^{Upa1}$ | pUL175 | Plasmid for the expression of PAM2L2 motif of Upa1 (DEDAADDDD**A**EFIYPNSY) carrying a point mutation (Aspartate 953 to Alanine) fused to an N-terminal 6xHis-Sumo tag. |
| pGX-G-Rrm4-M3-7H-H729A | pUL198 | Plasmid for the expression of G-Rrm4-M3-7H. MLLE3 domain of Rrm4 comprising amino acid residues from 679 to 792 with a point mutation (Histidine 729 to Alanine) fused to an N-terminal GST- tag. |
| pGX-G-Rrm4-M3-7H-R744A | pUL200 | Plasmid for the expression of G-Rrm4-M3-7H. MLLE3 domain of Rrm4 comprising amino acid residues from 679 to 792 with a point mutation (Arginine 744 to Alanine) fused to an N-terminal GST- tag. |
| pGX-G-Rrm4-M3-7H-I756G | pUL201 | Plasmid for the expression of G-Rrm4-M3-7H. MLLE3 domain of Rrm4 comprising amino acid residues from 679 to 792 with a point mutation (Isoleucine 756 to Glycine) fused to an N-terminal GST- tag. |
| pET28-HS-PAM2-L132A$^{Upa1}$ | pUL202 | Plasmid for the expression of the PAM2 motif of Upa1 (SQST**A**SPNASVFKPSRS) carrying a point mutation (Leucine 132 to Alanine) fused to an N-terminal 6xHis-Sumo tag. |
| pET28-HS-PAM2$^{FKYP}$ | pUL203 | Plasmid for the expression of the modified PAM2 motif of Upa1, FK**Y**P (SQSTLSPNASVFKYPSRS) fused to an N-terminal 6xHis-Sumo tag. |
| pET28-HS-PAM2L$^{Mkrn1_Hs}$ | pUL205 | Plasmid for the expression of the PAM2L motif of Human Mkrn1 (KSCPECRITSNFVIPSEY) fused to an N-terminal 6xHis-Sumo tag. |
| pGX-G-Rrm4-M3-7H-Q733A | pUL206 | Plasmid for the expression of G-Rrm4-M3-7H. MLLE3 domain of Rrm4 comprising amino acid residues from 679 to 792 with a point mutation (Glutamine 733 to Alanine) fused to an N-terminal GST- tag. |

| | | |
|---|---|---|
| pGX-G-Rrm4-M3-7H-F740A | pUL207 | Plasmid for the expression of G-Rrm4-M3-7H. MLLE3 domain of Rrm4 comprising amino acid residues from 679 to 792 with a point mutation (Phenylalanine 740 to Alanine) was fused to an N-terminal GST- tag. |
| pET28-HS-PAM2L2-A948L$^{Upa1}$ | pUL211 | Plasmid for the expression of the PAM2L2 motif of Upa1 (DEDALDDDDDEFIYPNSY) carrying a point mutation (Alanine 948 to Lucine) fused to an N-terminal 6xHis-Sumo tag. |
| pET28-HS-PAM2L$^{Taf7_Um}$ | pUL224 | Plasmid for the expression of the PAM2L motif of Ustilago Taf7 (GGGGKGFNIDDFIYPHGI) fused to an N-terminal 6xHis-Sumo tag. |
| pET28-HS-PAM2L1$^{Vps8_Um}$ | pUL225 | Plasmid for the expression of the PAM2L1 motif of Ustilago Vps8 (HQDDDNDNDDDFVYDGID) fused to an N-terminal 6xHis-Sumo tag. |
| pET28-HS-PAM2L2$^{Vps8_Um}$ | pUL226 | Plasmid for the expression of the PAM2L2 motif of Ustilago Vps8 (TDQAPSDDSFSFRYPHPL) fused to an N-terminal 6xHis-Sumo tag. |
| pGX-G-Ubr5-M_Hs | pUL234 | Plasmid for the expression of G-Ubr5-M. MLLE domain of Human Ubr5 comprising amino acid residues from 2389 to 2462, fused to an N-terminal GST- tag. |
| pGX-G-PABC1-M_Hs | pUL235 | Plasmid for the expression of G-PABC1-M. MLLE domain of Human PABC1 comprising amino acid residues from 545 to 636, fused to an N-terminal GST-tag |
| pGX-G-Rrm4-M3_Ri | pUL351 | Plasmid for the expression of G-Rrm4-M3_Ri. MLLE3 domain of Rhizophagus irregularis Rrm4 comprising amino acid residues from 557 to 639, fused to an N-terminal GST-tag |

#### Table S12: DNA oligonucleotides used in this study

| Designation | Nucleotide sequence (5' ➔ 3') | Construct generated |
|---|---|---|
| RL1456 | ATAGGATCCCATATGGTGGCCATTACGGCCAGT | pUMa2187 |
| RL1457 | CTAGGAATTCTCACGCGTTGGCCTTAGC | pUMa2187 |
| AB45 | CGGCCATATGGGCAGCAGCCATCATC | HS-PAM2/L (kanR)_FP |
| AB46 | CTCACTCGAGTTAGGATCGGGACGGCTTGAAGACGGAGGCGTTGGGAGACAAGGT GCTTTGCGAACCACCAATCTGTTCTCTGTGAGC | pUMa4296 |
| AB47 | CTCACTCGAGTTAGTCGGCTCCTGGGTAGACAAAGTCATCTTGATCTTCCTCTTGGT CTGCAGCCTCACCACCAATCTGTTCTCTGTGAG | pUMa4297 |
| AB48 | CTCACTCGAGTTAGTACGAGTTCGGGTAGATGAATTCATCGTCATCGTCATCGGCCG CATCCTCGTCACCACCAATCTGTTCTCTGTGAG | pUMa4298 |
| MB696 | TCGACTCGAGTCACTTGTTCAGACCTGCAGC | pUMa4701,pUMa4704 |
| CD503 | CATGCCATGGGAGCTAGCGCGGCCGCCAGCTCCGGTCTGCCTCTCC | pUMa4701 |
| CD605 | ATGCATCATATGCATCATCATCATCATCACACATTATCCACGCTTGCTGC | pUMa4704 |
| CD694 | ACTGCACATATGCATCATCATCATCATCACAGTCCCGAGGAGCAGAAGC | pUMa4794 |
| AB611 | ATGCGAATTCGGTACCTTACGCAGAGTCGTTCTGTTG | pUMa4794 |
| CD658 | CTTTAAGAAGGAGATATACATATGGGCAGCAGCCATCATCATC | HS-PAM2/L (ampR)_FP |
| CD700 | GTGGTGGTGGTGGTGCTCGAGTTACATGTTCGGAAAAATAAATTCTTTCGCGTTCGG GCTCAGCGCGCTACCACCAATCTGTTCTCTGTG | pUL123 |
| CD657 | GCAGCCGGATCTCACTCGAGTTAAAAGCCGCGTTTGCCCGGCTGATAAAACAGCGG CGCGTTATCATCGCCATCATCGGTATCACCACCAATCTGTTCTCTG | pUL124 |
| CD701 | GAGGTTCCATGGCACATATGACATTATCCACGCTTGCTGC | GST-M3_FP/ pUL130 |
| CD702 | CTAGGCCGCGTTGGCCTCGAGTCACTTGTTCAGACCTGCAGC | GST-M3_RP/ pUL130 |
| EF034 | ACCCAGCTTCGCCTTCTGATCGTGCGCTGCTTTGCC | pUL131/ pUL206/ pUL220/ pUL222 |
| EF009 | CTAGGCCGCGTTGGCCTCGAGTCACTTGTTCAGACC | GST-M3_RP |
| EF169 | CGATCAGAAGGCGAAGCTGGGTGATCAGCTCGCCAAAAAGATCCGTACGTTCGGCG TC | pUL131 |
| EF534 | GGTGGTGGTGGTGCTCGAGTTAAAACGGCTGCGGGGTCACAAAGGTAAAATCGCTC GGGGTGCGGCAGGTCGGGCAACCACCAATCTGTTCTCTG | pUL132 |
| EF535 | GGTGGTGGTGGTGGTGCTCGAGTTACGGCTGGCCCGGCACAAATTCAATCGCGTTC ACCCAATCTTCGCTGCCCGCACCACCAATCTGTTCTCTG | pUL133 |

| | | |
|---|---|---|
| EF536 | GGTGGTGGTGGTGGTGCTCGAGTTAGCGCGCCACCGGCTGAAACGCCTGCACGCTC GCGCTCAGGTTGCTGCGGCTACCACCAATCTGTTCTCTG | pUL134 |
| CD695 | GTTAGCAGCCGGATCTCACTCGAGTTAGTCGGCTCCTGGGTAGACAAAGTCATCAC CACCAATCTGTTCTCTGTG | pUL138 |
| CD696 | GTTAGCAGCCGGATCTCACTCGAGTTAGTACGAGTTCGGGTAGATGAATTCATCAC CACCAATCTGTTCTCTGTG | pUL139 |
| CD710 | GTGGTGGTGGTGGTGCTCGAGTTACTGCAGGCCTTTCCACGGCACGCCCGGATGAA ATTCCGGCGGCCAGTTACCACCAATCTGTTCTCTG | pUL146 |
| CD759 | GGTGGTGGTGGTGCTCGAGTTAGTAAGAGTTCGGGTAGATGAATTCGTCCGCGTTC GGAGACAGGGTAGACTGAGAACCACCAATCTGTTCTCTG | pUL157 |
| CD769 | GGTGGTGGTGGTGGTGCTCGAGTTATTTAACACCCGGAACGAATTCTTTCGCGTTCG GGTTCAGGTTAGATTTAACACCACCAATCTGTTCTCTGTG | pUL162 |
| CD808 | GTGGTGGTGGTGGTGCTCGAGTTAATCCGCGCCCGGCGCCACAAAATCATCCTGAT CTTCTTCCTGATCCGCCGCTTCACCACCAATCTGTTCTCTG | pUL166 |
| CD809 | GTGGTGGTGGTGGTGCTCGAGTTAATAGCTGTTCGGCGCAATAAATTCATCATCATC ATCATCCGCCGCATCTTCATCACCACCAATCTGTTCTCTG | pUL167 |
| CD810 | GTGGTGGTGGTGGTGCTCGAGTTAGCTGCGGCTCGGTTTAAACACGCTATCATCATC ATCCGCCGCATCTTCATCACCACCAATCTGTTCTCTG | pUL168 |
| CD811 | GTGGTGGTGGTGGTGCTCGAGTTAATCCGCGCCCGGATACACCGCATCATCCTGAT CTTCTTCCTGATCCGCCGCTTCACCACCAATCTGTTCTCTG | pUL169 |
| CD812 | GTGGTGGTGGTGGTGCTCGAGTTAATAGCTGTTCGGATAAATCGCTTCATCATCATC ATCATCCGCCGCATCTTCATCACCACCAATCTGTTCTCTG | pUL170 |
| CD813 | GTGGTGGTGGTGGTGCTCGAGTTAATCCGCGCCCGCATACACAAAATCATCCTGAT CTTCTTCCTGATCCGCCGCTTCACCACCAATCTGTTCTCTG | pUL171 |
| CD814 | GTGGTGGTGGTGGTGCTCGAGTTAATAGCTGTTCGCATAAATAAATTCATCATCATC ATCATCCGCCGCATCTTCATCACCACCAATCTGTTCTCTG | pUL172 |
| CD815 | GTGGTGGTGGTGGTGCTCGAGTTAGCTGCGGCTCGGTTTCGCCACGCTCGCGTTCGG GCTCAGGGTGCTCTGGCTACCACCAATCTGTTCTCTG | pUL173 |
| CD816 | GTGGTGGTGGTGGTGCTCGAGTTAATCCGCGCCCGGATACACAAAATCCGCCTGAT CTTCTTCCTGATCCGCCGCTTCACCACCAATCTGTTCTCTG | pUL174 |
| CD817 | GTGGTGGTGGTGGTGCTCGAGTTAATAGCTGTTCGGATAAATAAATTCCGCATCATC ATCATCCGCCGCATCTTCATCACCACCAATCTGTTCTCTG | pUL175 |
| EF001 | GCAAAGCAGCGGCGGATCAGAAGCAGAAGCTGGGTGATCAGCTC | pUL198 |
| EF008 | CTGCTTCTGATCCGCCGCTGCTTTGCCCTGAAGCGAGTCG | pUL198 |
| EF013 | CTTCAAAAAGATCGCTACGTTCGGCGTCAAAGGCGCACCCAAGC | pUL200 |
| EF014 | CGAACGTAGCGATCTTTTTGAAGAGCTGATCACCCAGC | pUL200 |
| EF015 | GCACCCAAGCTCACCGGCCACTTGTTGGATTCCGAAGACTTGCGGGCG | pUL201 |

| | | |
|---|---|---|
| EF016 | CCAACAAGTGGCCGGTGAGCTTGGGTGCGCCTTTGACGCC | pUL201 |
| EF017 | GTTAGCAGCCGGATCTCACTCGAGTTAGGATCGGGACGGCTTGAAGACGGAGGCGTTGGGAGACGCGGTGCTTTGCGAACCACCAATCTGTTCTCTGTG | pUL202 |
| EF018 | AGCAGCCGGATCTCACTCGAGTTAGGATCGGGACGGGTACTTGAAGACGGAGGCGTTGGGAGACAAGGTGCTTTGCGAACCACCAATCTGTTCTCTGTG | pUL203 |
| EF020 | AGCCGGATCTCACTCGAGTTAATATTCGCTCGGAATCACAAAGTTGCTGGTAATGCGGCATTCCGGGCAGCTTTTACCACCAATCTGTTCTCTGTGAGC | pUL205 |
| EF033 | GCAGCGCACGATCAGAAGGCGAAGCTGGGTGATCAGCTCTTCAAAAAGATCC | pUL206/ pUL222 |
| EF035 | CAGAAGCTGGGTGATCAGCTCGCCAAAAAGATCCGTACGTTCGGCGTCAAAGGC | pUL207 |
| EF036 | ATCTTTTTGGCGAGCTGATCACCCAGCTTCTGCTTCTGATCG | pUL207 |
| EF532 | GGTGGTGGTGGTGCTCGAGTTAATAGCTGTTCGGATAAATAAATTCATCATCATCATCATCTAACGCATCTTCATCACCACCAATCTGTTCTCTG | pUL211 |
| EF162 | CCACATAATAAGCACAAGCTCGAGGCTGCTGAGGCTTACACATTATCC | pUL221 |
| EF163 | GGATAATGTGTAAGCCTCAGCAGCCTCGAGCTTGTGCTTATTATGTGG | pUL221 |
| EF164 | CAGAAGGAGGGTGACATGCCCTAGGTGCGCTGAACTCTCCACCGC | pUL221 |
| EF165 | ACTACGCCCTGGAAGTACTGTTCCAGGGGCCCCTCCCTAAAGCGGATCCAGCGACGG | pUL219 |
| EF166 | CAGTACTTCCAGGGCGTAGTCGGGCACGTCGTAAGGGTAGTAAGCCTCAGCAGCGCCAGG | pUL219 |
| EF167 | GCCAGCGGTCTTGTAGACGAAGAGCAATTGCGCTCGCTCAGCACTGTCG | pUL219/ pUL220<br>pUL222 |
| EF169 | CGATCAGAAGGCGAAGCTGGGTGATCAGCTCGCCAAAAAGATCCGTACGTTCGGCGTC | pUL220 |
| AB351 | CCACGGCGCGCCACCGGTTTAGCGGTGACCGAGTTTCGAG | pUL219/ pUL220<br>pUL221/ pUL222 |
| EF87 | GTGGTGGTGGTGGTGCTCGAGTTAGATACCGTGCGGGTAGATGAAGTCGTCGATGTTGAAACCTTTACCACCACCACCACCACCAATCTGTTCTCTGTG | pUL224 |
| EF92 | GTGGTGGTGGTGGTGCTCGAGTTAGTCGATACCGTCGTAAACGAAGTCGTCGTCGTTGTCGTTGTCGTCGTCCTGGTGACCACCAATCTGTTCTCTGTG | pUL225 |
| EF107 | GTGGTGGTGGTGGTGCTCGAGTTACAGCGGGTGCGGGTAACGGAAAGAGAAAGAGTCGTCAGACGGCGCCTGGTCGGTACCACCAATCTGTTCTCTGTG | pUL226 |
| EF944 | GGGAGGTTCCATGGCACATATGCCCGAGTACAAGGAGATCGAGGAG | pUL351 |
| EF945 | CGTTGGCCTCGAGTCACTTGGGCACAACCTTCTCTGC | pUL351 |
| EF395 | TGAACGATCGCCGGGCGGCCGGCGCGCCTTAGCGGTGACCGAGTTTCGAG | pUL254 |
| EF231 | GCTGGAATTCGCCCTTTAGCCATGTCCGATTCGATTTACGC | pUL254 |
